# Mechanical model of muscle contraction. 1. Force-velocity relationship

**DOI:** 10.1101/2019.12.16.878793

**Authors:** S. Louvet

## Abstract

The two parameters that determine the functionality of a skeletal muscle fiber are the tension (T) exerted at its two endpoints and the shortening speed (V), two mechanical characteristics. We established a relationship between T and V by developing a theoretical model of muscle contraction based on the swinging lever arm hypothesis. At the nanoscale, force and movement are generated by the myosin II heads during the working stroke (WS). The change in conformation of a myosin head during the WS is characterized by the rotation of the lever correlated to the linear displacement of the motor domain. The position of the lever is marked by the angle θ. The maximum variation of θ between the two limits θ_up_ and θ_down_ relating to the two positions *up* and *down* is usually given equal to 70°. When the angle θ is between θ_up_ and θ_down_, the WS is triggered in three modes, fast, slow or very slow. During the isometric tetanus plateau, θ is uniformly distributed between the two angles θup and θ_T_ separated by a usual difference of 50°. Consequently during isometric tetanus plateau there is a 20° interval between θ_T_ and θ_down_ where no head is found in WS. We link this absence to the slow detachment of the heads whose orientation of the levers is between θ_T_ and θ_down_ during the rise to the isometric tetanus plateau. The equation between T and V refers to these four occurrences: fast, slow or very slow initiations of the WS between θ_up_ and θ_down_, then slow detachment between θ_T_ and θ_down_. The equation is constructed from the geometric data of the myosin head and the time constants of the cross-bridge cycle reactions associated with these four events. The biphasic aspect of the curve is explained by the slow detachment that occurs only at very slow speeds. An additional term, derived from the viscosity present as soon as the velocity increases completes the equation. An adequate fit between the model and examples from the physiological literature is found (r^2^ > 99%).

## Introduction

The relationship between tension and shortening velocity has been studied in skeletal fiber for almost a century. Several models have been proposed, and among them Hill’s hyperbolic model [1] remains the reference. These empirical models are predictive but do not include any data related to the myosin II head, the nanomotor responsible for the motricity of a half-sarcomere (hs), a myofibril and the fiber. Our objective is to present an equation that combines the mechanical characteristics of the myosin head and the time characteristics of the cross-bridge cycle composed of a sequence of mechanical-chemical reactions preceding and interrupting the WS [2,3].

The relationship between T and V is delivered without preamble because its demonstration requires five other articles. Based on the postulate of swinging lever arm [3,4], accompanying Paper 2 analyses the geometry and kinetics of a WS myosin head. The conclusions of Paper 2 allow the study of the distribution of the θ orientation of levers (S1b) belonging to the WS heads (Fig 1a). Conducted in Paper 3, this work leads to a uniform density for θ, especially between θ_up_ and θ_T_ during the isometric tetanus plateau (Fig 1b). With the results of Papers 2 and 3, accompanying Paper 4 determines the tension of the isometric tetanus plateau (T0) and the minimum tension (T1) reached after a sudden shortening of the fiber by a length step. Paper 5 calculates the rise tension from T1 to T0 after the phase 1 of a length step. Paper 6 studies staircase shortening, i.e. the shortening of a succession of length steps. By gradually decreasing the size and duration of the step, the staircase shortening tends towards a shortening at a constant speed. The precepts of infinitesimal calculation and integral calculation provide the equation between T and V. The determination of the tension T1 in Paper 4 reveals the presence of viscosity during phase 1 of a length step, i.e. when shortening at a constant but variable speed from one step to the next. This result leads to the calculation of the viscosity force (T_visc_) as a function of the hs shortening speed. The addition of the term Tvisc completes the general wording of the modelling between T and V.

**Fig 1.**
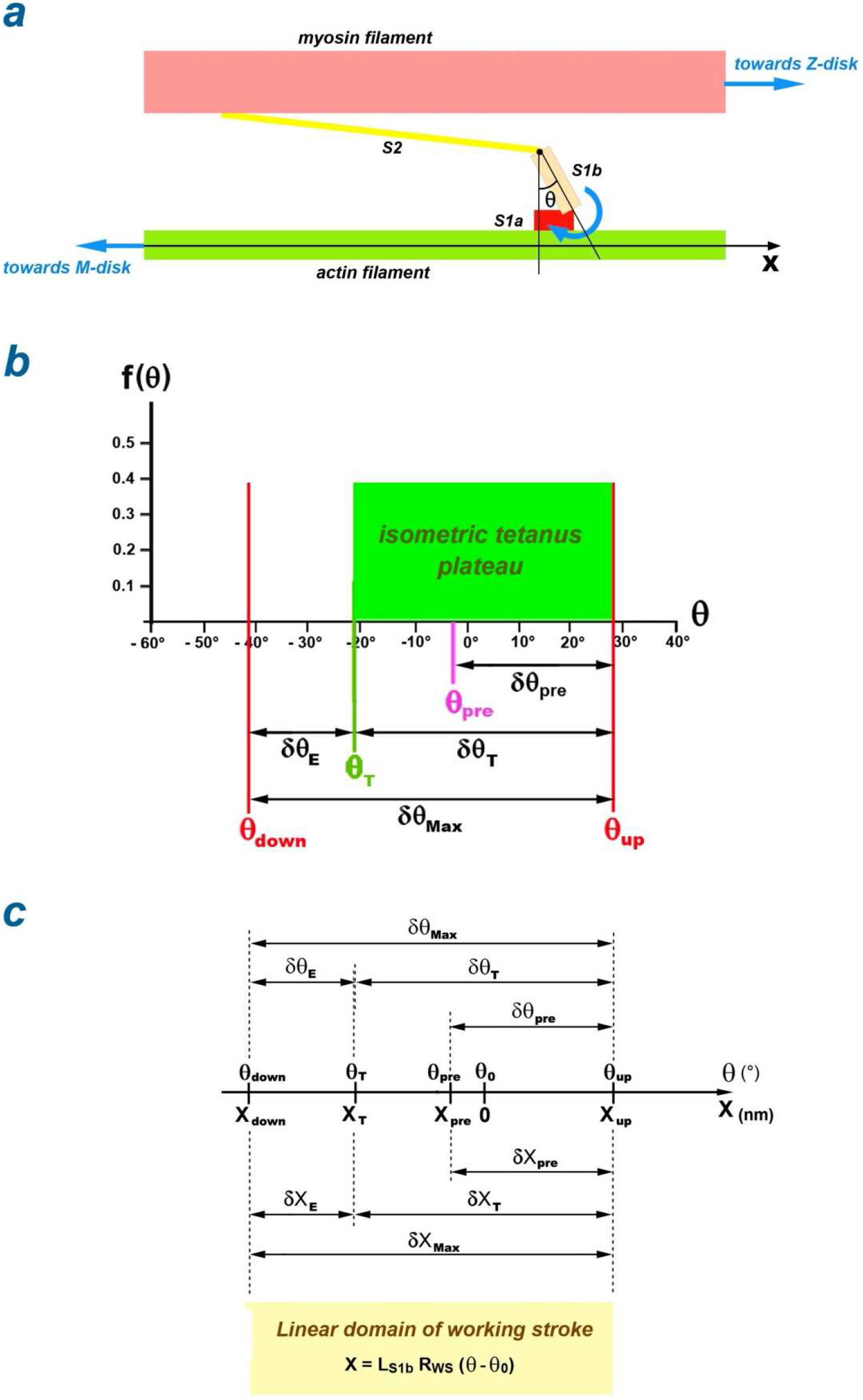
Geometric variables related to a myosin II head. (a) Definition of the angle θ characterizing the position of lever S1b of a WS myosin head in a half-sarcomere on the right. (b) Uniform distribution of θ, represented by a green rectangle bounded by θ_T_ and θ_up_ in a half-sarcomere on the right during isometric tetanus plateau. (c) Affine relationship between θ and X, the abscissa of the myosin-binding site on actin along the OX axis with graphic correspondence between the linear intervals, δX_Max_, δX_T_, δX_E_, δX_pre_, and the angular intervals, δθ_Max_, δθ_T_, δθ_E_, δθ_pre_.

Our model produces curves that are consistent with those presented in the articles published since 1938 and correctly responds to experimental variants such as initial sarcomere length, fiber length, inter-filament distance and temperature.

## Methods

### Linear domain specific to the Working Stroke

In accompanying Paper 2, the values of the two angles θ_up_ and θ_down_ are determined. These two limits define a linear interval (Fig 1c) for the abscissa of the motor domain of the myosin head correlated to the angular position of the lever (Fig 1a), such that:
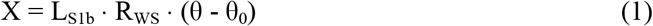

where X is the abscissa of the myosin-binding site on the actin molecule belonging to the actin filament customizing the longitudinal axis (OX); θ is the angle that the lever (S1b) forms with the axis perpendicular to OX in the fixed plane where S1b moves; L_S1b_ is the length of S1b varying between 8 and 10 nm; R_WS_ is a dimensionless data characteristic of the geometry of the myosin head evaluated at 0.95, approximately; θ_0_ is the middle of δθ_T_ (Fig 1c).

With the affine function defined in (1), the two angular ranges δθ_Max_ and δθ_T_ presented in the abstract correspond to the two linear ranges δX_Max_ and δX_T_ (Figs 1b and 1c). Before the length step, if the strongly linked heads likely to initiate a WS were already in WS, the θ angle of their levers would have a value higher (lower) than θ_up_ in a hs on the right (left) and would be distributed uniformly over the angular range δθ_pre_. After the length step, the θ angle of the head levers quickly initiating a WS is distributed between θ_pre_ and θ_up_ over an angular range equal to δθ_pre_ by symmetry with respect to θ_up_, located within the linear domain and consequently associated with the linear range δX_pre_ (Fig 1c).

The X_down_, X_up_ and X_T_ abscissa correspond to the angles θ_down_, θ_up_ and θ_T_ (Fig 1c). Their presence in some of the following equations requires calculation:

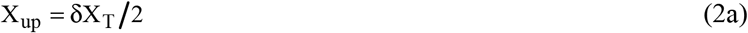

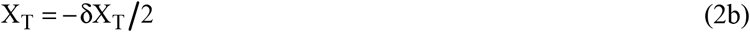

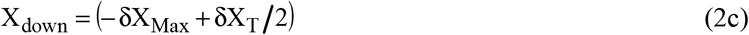

### Force-velocity relation

The relative force or tension (pT) and the module of the hs shortening speed (u) are defined:

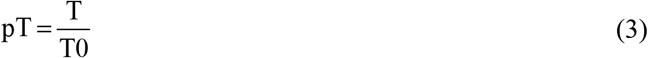

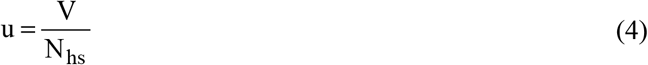

where T0 is the constant tension of the isometric tetanus plateau preceding the different force steps leading to the points of the T/V relationship; N_hs_ is the number of hs per myofibril.

With (3) and (4), it is equivalent to searching for a relationship between T and V or between pT and u. We pose:

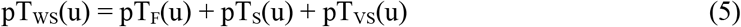

where pT_WS_ is the relative tension generated by all WS heads in a hs that shortens to velocity u; pT_F_, pT_S_ and pT_VS_ are the relative tensions generated by WS heads that initiated their WS quickly, slowly or very slowly, respectively.

The events leading to the initiation of a WS in the 3 exclusive modes, fast, slow or very slow, are presented in paragraph B.3 of Supplement S1.B.

We introduce:

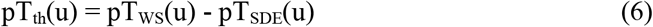

where pT_th_ (th for *theoretical*) is the relative tension with consideration of slowly detaching heads; pT_SDE_ is the relative tension generated by the slow detachment of the myosin heads previously in WS.

The event relating to the slow detachment is discussed in paragraph B.7 of Supplement S1.B.

The 4 terms pT_F_, pT_S_, pT_VS_ and pT_SDE_ present in equations (5) and (6) are established in Supplement S6.L of accompanying Paper 6 and their respective equations are delivered below.

#### 1/ Fast initiation

The event where a myosin head quickly initiates a WS is called as {startF}; F for *Fast*. The relative tension (pT_F_) associated with {startF} is written as a function of u:

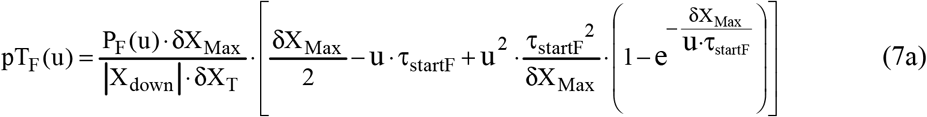

where P_F_ is the proportion of heads starting quickly a WS, a function of u, such that:

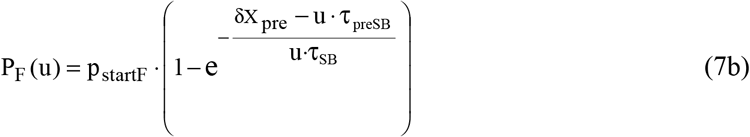

where p_startF_ is the maximum proportion of heads that quickly initiated a WS during rise to the isometric tetanus plateau; τ_startF_ is the time constant of the event {startF}; τ_preSB_ and τ_SB_ are the occurrence delay and time constant of the strong binding event noted {SB}; δX_Max_, δX_T_ and δX_pre_ are three linear ranges that were presented in the previous paragraph (Fig 1c); X_down_ is an abscissa defined in (2c).

The {SB} event leading to the strong binding state is discussed in paragraph B.4 of Supplement S1.B.

#### 2/ Slow initiation

The event where a head slowly initiates a WS is called as {startS}; S for *Slow*. The relative tension (pT_S_) associated with {startS} is formulated as a function of u:

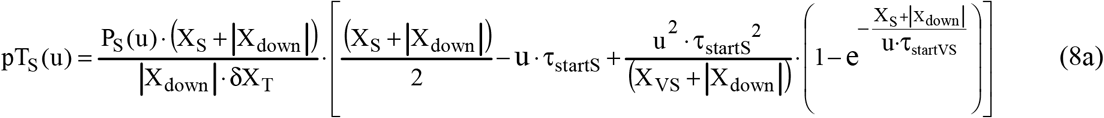

where P_S_ is the proportion of heads slowly starting a WS, a function of u, equal to:

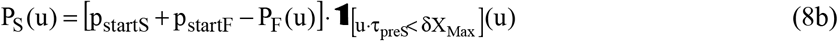

where X_S_ is an abscissa dependent on u in relation to the event {startS} such that:

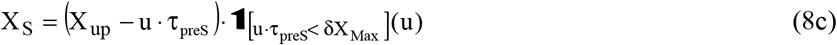

where p_startS_ is the maximum proportion of heads that slowly initiated a WS during the rise to the isometric tetanus plateau; X_up_ is determined in (2a); τ_preS_ and τ_startS_ are the occurrence delay and time constant of {startS}; **1** is the indicator function formulated in (A2b) in Supplement S1.A.

#### 3/ Very Slow initiation

The event where a head very slowly initiates a WS is called as {startVS}; VS for *Very Slow*. The relative tension (pT_VS_) associated with {startVS} is enounced as a function of u:

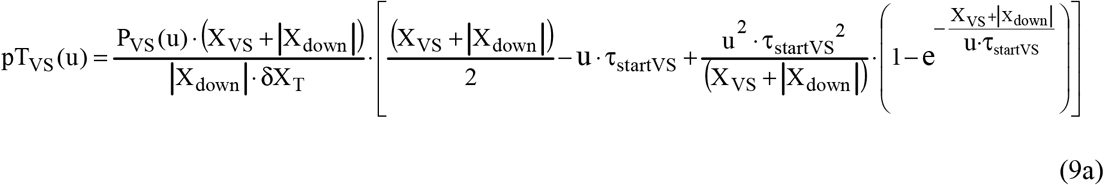

where P_VS_ is the maximum proportion of heads starting very slowly a WS during rise to the isometric tetanus plateau, a function of u, equal to:

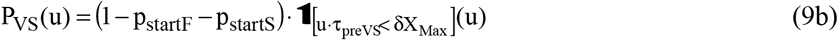

where X_VS_ is an abscissa dependent on u in relation to the event {startVS} such that:

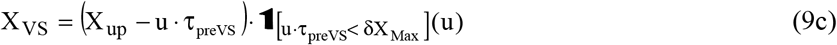

where τ_preVS_ and τ_startVS_ are the occurrence delay and time constant of {startVS).

#### 4/ Slow detachment

The event where a head previously in WS slowly detaches is called as {SlowDE}. The relative tension (pT_SDE_) associated with {SlowDE} is expressed as a function of u:

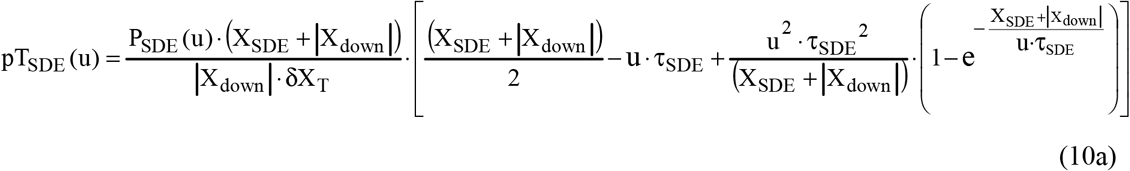

where P_SDE_ is the proportion of slowly detaching heads, a function of u, equal to:

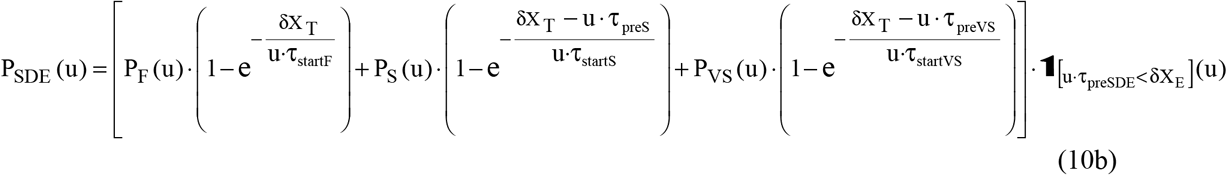

where X_SDE_ is an abscissa dependent on u in relation to the event {SlowDE} defined as:

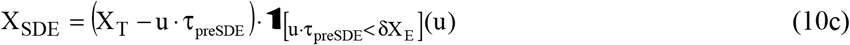

where X_T_ is determined in (2b); τ_preSDE_ and τ_SDE_ are the occurrence delay and time constant of {SlowDE).

All parameters in equations (7a) to (10c) are explained in Supplement S1.B and standard values are given in Table B1.

### Presence of viscosity

In sub-paragraph (J.16.5) of Supplement S4.J to accompanying Paper 4, it is established that the relative tension induced by viscosity (pT_Visc_) is a linear function of u:

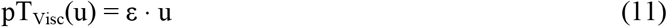

where ε is a proportionality coefficient that depends on the experimental methodology; u is expressed in modulus.

The parameter ε is calculated under standard conditions at a temperature of 1°C for a fast fiber isolated from the *tibialis anterior* muscle in two species of frogs:

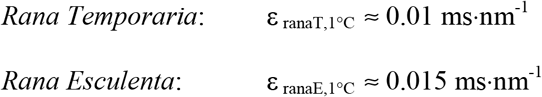

The method for determining these two data is used to evaluate the viscous coefficient ε when tests are performed with different procedures. Thus, an increase in temperature causes a decrease of ε, and a decrease in the inter-filament distance or lattice produces an increase of ε.

### Complete formulation of the relationship between tension and speed of shortening

With equations (5), (6) and (11), the relationship between pT and u is formulated:

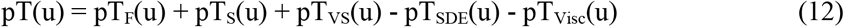

### Relationship between the number of heads in WS per hs and the speed of shortening

The total of the myosin heads in WS (Λ) contributing to the shortening at constant velocity of any hs of the muscle fiber is the sum of 4 countings related to the 4 events {startF}, {startS}, {startVS} and {SlowDE} such that:

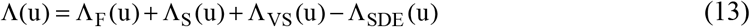

where Λ_F_, Λ_S_ and Λ_VS_ are the number of WS heads per hs after fast, slow or very slow WS initiation; Λ_SDE_ is the number of heads per hs that detach slowly.

The 4 terms of the right-hand member of the expression (13) are calculated in the Supplement S6.L of Paper 6 and are formulated:

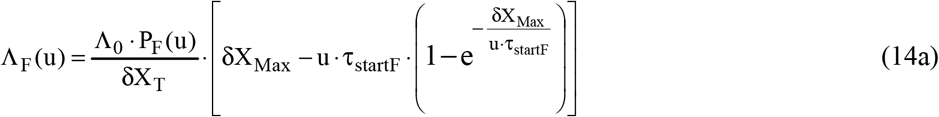

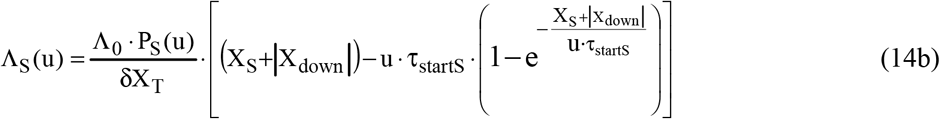

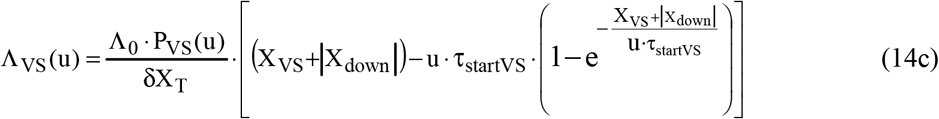

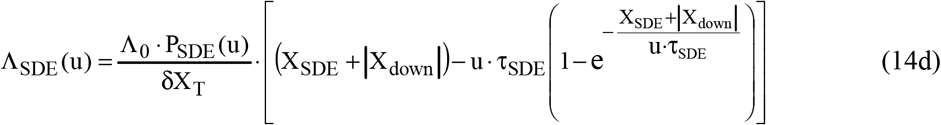

where Λ_0_ is the number of myosin heads in WS per hs during the isometric tetanus plateau.

### Maximum tensions

From expressions (5), (6) and (12), the maximum tensions pT0_WS_, pT0_th_ and pT0 reached during the isometric tetanus plateau, i.e. u=0, are equal to:

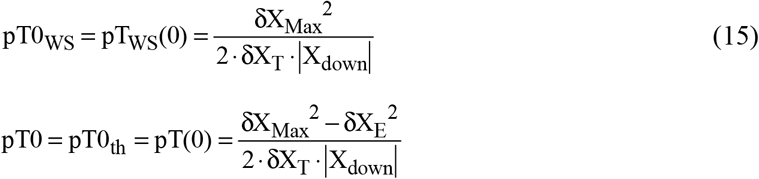

By squaring equality « δX_Max_ = δX_T_ + δX_E_ », we check with (2c):

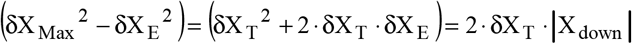

The equalities « pT0 = pT0_th_ = 1 » are corroborated and it implies:

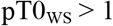

### Maximum shortening speeds

The theoretical maximum speed of a hs (u_Max,th_) corresponds to the limit condition of achievement of the {startF} event, i.e. according to (7b):

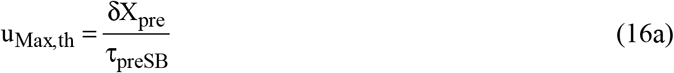

With (4), the theoretical maximum fiber velocity (V_Max,th_) is:

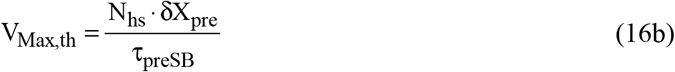

The actual maximum velocities uMax and VMax are lower than u_Max,th_ and V_Max,th_ respectively, because the forces due to viscosity slow down the shortening at high speeds.

### Objective

The purpose of the paper is to compare the previous equations with the experimental points of the Force/Velocity relationship drawn from examples in the physiological literature.

### Statistics

A linear regression is performed between the measured tension values (points recorded on the figures of the articles listed) and the theoretical tension values calculated at the same velocity. The regression line goes through the origin, such that:

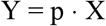

where Y characterizes the values of the theoretical tensions and X those of the experimental tensions; p is the slope of the regression line equal to:

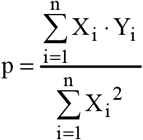

The determination coefficient is defining as:

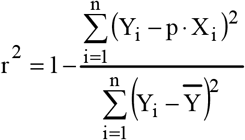

### Algorithmic

The equations are programmed (Visual Basic 6 language) to obtain the plot of the relationships of u or V according to T, T_th_, TW_S_, pT, pT_th_ or pT_WS_.

With pΛ=Λ/Λ_0_, the relationship pΛ/pT_th_ is derived from equations (13) and (6), and the relationship pΛ/pT of (13) and (12).

### Adequacy between experimental points and theoretical graphical representation

The adjustment is made visually with help of the trial and error method, by searching for the p and r^2^ values closest to 1, and by allocating to the theoretical parameters values compatible with those in the literature, on the one hand, and with those credited in the calculations of accompanying Papers 2 to 6, on the other hand.

## Results

### Reference plot (Fig 2 associated with the “REF” column in Table 1)

K.A.P. Edman has particularly studied the Force/Velocity relation of muscle fibers in *Rana Temporaria* by highlighting the biphasic shape of the curve [5,6,7]. We have chosen the points in Fig 2A in [8] as a reference for the F-V relationship. We remind that the force or tension T is calculated by taking into account the three WS initiation events and the slow detachment event in the presence of viscosity, that the tension T_th_ is evaluated with the 4 events in the absence of viscosity and that the tension TWS is determined with the three WS initiation events only.

**Fig 2.**
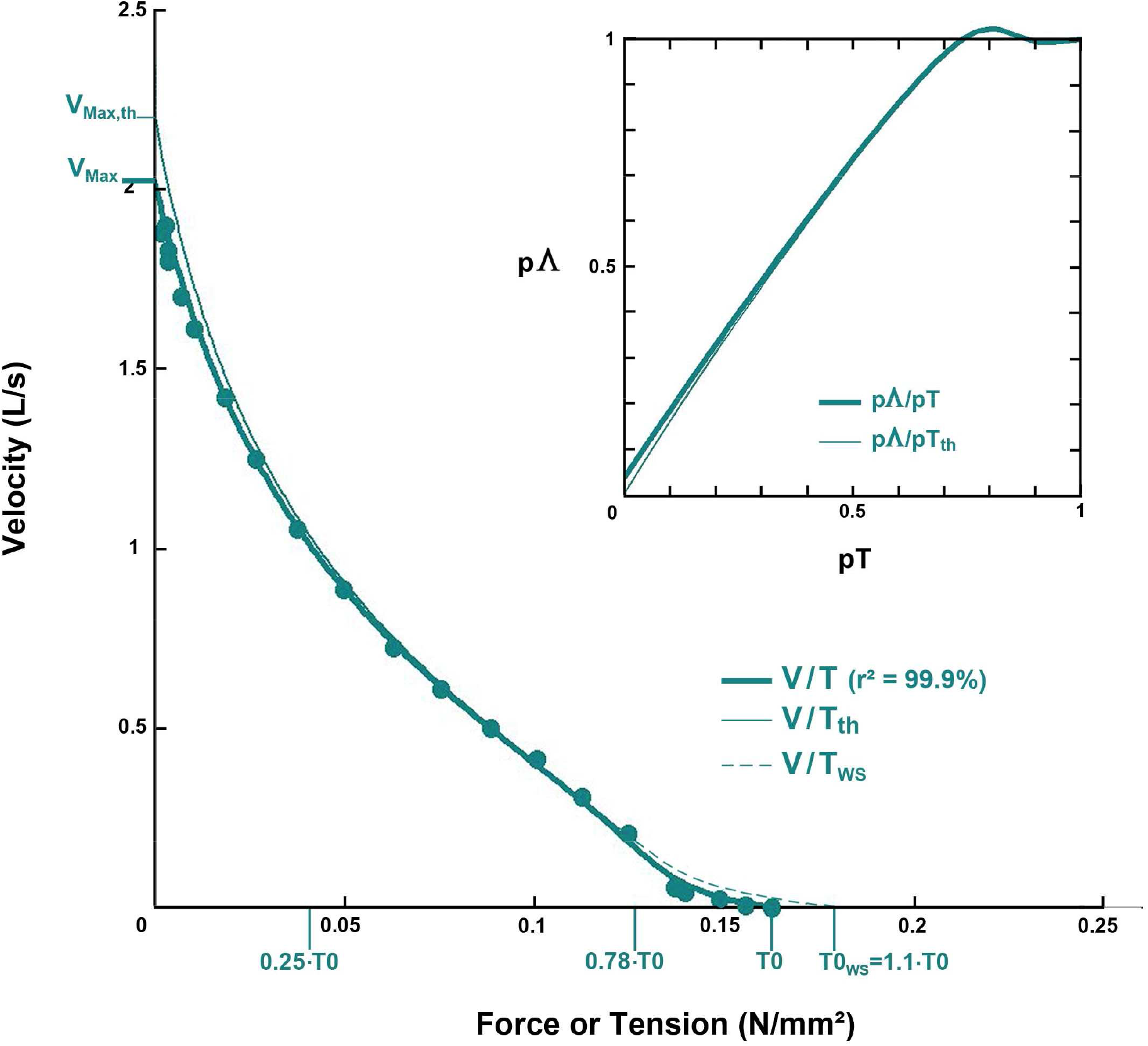
Reference plots of the Force-Velocity relationship. L is a unit referring to the initial length of the fiber tested (L0), i.e. V measured in mm/s is divided by L0. The relationships V/T, V/T_th_ and V/T_WS_ are plotted with a thick solid line, a fine solid line and dotted lines, respectively, according to equations (12), (6) and (5). The points are from Fig 2A in [8]. Inset: the relative number of WS heads (pΛ) is plotted as a function of the relative tensions pT and pT_th_, represented by thick and fine solid lines, respectively.

**Table 1.** Values of the characteristic parameters of a muscle fiber, a half-sarcomere and a myosin head isolated from the *tibialis anterior* muscle of frog *(Rana Temporaria*) relative to the plots in Figs 2 and 3.

| | REF | 1.85 $\mu\text{m}$ | 2.1 $\mu\text{m}$ | 2.3 $\mu\text{m}$ | 2.6 $\mu\text{m}$ |
| --- | --- | --- | --- | --- | --- |
| $\Gamma$ | 1°C | 1°C | 1°C | 1°C | 1°C |
| T0 | 163 kPa | 246 kPa | 257 kPa | 237 kPa | 186 kPa |
| L0 | 6.3 mm | 7.4 mm | 8.35 mm | 9.15 mm | 10.35 mm |
| N<sub>hs</sub> | 6000 | 8000 | 8000 | 8000 | 8000 |
| L0<sub>s</sub> | 2.1 $\mu\text{m}$ | 1.85 $\mu\text{m}$ | 2.1 $\mu\text{m}$ | 2.3 $\mu\text{m}$ | 2.6 $\mu\text{m}$ |
| p<sub>startF</sub> | 0.6 | 0.63 | 0.62 | 0.61 | 0.57 |
| p<sub>startS</sub> | 0.3 | 0.25 | 0.28 | 0.27 | 0.28 |
| p<sub>startVS</sub> | 0.1 | 0.12 | 0.1 | 0.12 | 0.15 |
| $\varepsilon$ | 0.01 ms·nm<sup>-1</sup> | 0.01 ms·nm<sup>-1</sup> | 0.01 ms·nm<sup>-1</sup> | 0.01 ms·nm<sup>-1</sup> | 0.01 ms·nm<sup>-1</sup> |
| u<sub>Max,th</sub> | 2.31 nm·ms<sup>-1</sup> | 2.4 nm·ms<sup>-1</sup> | 2.4 nm·ms<sup>-1</sup> | 2.4 nm·ms<sup>-1</sup> | 2.4 nm·ms<sup>-1</sup> |
| u<sub>Max</sub> | 2.14 nm·ms<sup>-1</sup> | 2.12 nm·ms<sup>-1</sup> | 2.11 nm·ms<sup>-1</sup> | 2.04 nm·ms<sup>-1</sup> | 2.15 nm·ms<sup>-1</sup> |
| $\delta X_{\text{Max}}$ | 11.5 nm | 11.5 nm | 11.5 nm | 11.5 nm | 11.5 nm |
| $\delta X_{\text{T}}$ | 8 nm | 7 nm | 8 nm | 8 nm | 8 nm |
| $\delta X_{\text{pre}}$ | 6 nm | 6 nm | 6 nm | 6 nm | 6 nm |
| $\tau_{\text{preSB}}$ | 2.6 ms | 2.5 ms | 2.5 ms | 2.5 ms | 2.5 ms |
| $\tau_{\text{SB}}$ | 6 ms | 8 ms | 6.5 ms | 7 ms | 7.5 ms |
| $\tau_{\text{startF}}$ | 0.7 ms | 0.7 ms | 0.7 ms | 0.7 ms | 0.7 ms |
| $\tau_{\text{preS}}$ | 3.5 ms | 3.5 ms | 3.5 ms | 3.5 ms | 3.5 ms |
| $\tau_{\text{startS}}$ | 25 ms | 25 ms | 25 ms | 25 ms | 25 ms |
| $\tau_{\text{preVS}}$ | 80 ms | 80 ms | 80 ms | 80 ms | 80 ms |
| $\tau_{\text{startVS}}$ | 90 ms | 90 ms | 90 ms | 90 ms | 90 ms |
| $\tau_{\text{preSDE}}$ | 5 ms | 5 ms | 5 ms | 5 ms | 5 ms |
| $\tau_{\text{SDE}}$ | 18 ms | 18 ms | 18 ms | 18 ms | 18 ms |
| R<sup>2</sup> | 99.9% | 99.9% | 99.9% | 99.9% | 99.9% |

The calculations are made using the data in the “REF” column of Table 1 and a good agreement is obtained for the V/T relationship with r^2^=99.9% (Fig 2; thick blue line).

For T ≥ 0.78·T0, the convexity of V/T is more pronounced than that of V/T_WS_ (Fig 2; blue dotted line), affirming the characteristic biphasic aspect of the V/T curve. The value of T0_WS_ is determined according to (15). The difference between the two curves underlines the role of slow detachment at low speeds.

A difference is observed between the V/T and V/T_th_ plots (Fig 2; thin blue line) for tension values below 25% of T0. The presence of viscosity is quantified using the coefficient ε introduced in equation (11). There is a 7% difference between V_Max_ and V_Max,th_: the influence of viscosity is minimal but not negligible.

In the inset, the number of myosin heads in WS is approximately stable for high tensions (pT > 70%) then a quasi linear decrease is observed from (0.7·T0) to very low tension values.

where the first part is related to the fiber: Γ is the experimental temperature; T0 is the tension of the tetanus plateau in isometric conditions; L0 is the initial length of the fiber.

The 2^nd^ part is related to the sarcomere and the half-sarcomere (hs): N_hs_ is the number of hs per myofibril; L0_s_ is the initial length of the sarcomere; p_startF_, p_startS_ and p_startVS_ are the percentage of WS heads per hs during an isometric tetanus plateau according to the WS initiation mode, fast, slow and very slow; ε is the proportionality coefficient between the relative viscous tension (pT_Visc_) and the hs shortening velocity (u); u_Max,th_ is the hs theoretical maximum velocity without viscosity equal to δX_pre_/τ_preSB_; u_Max_ is the hs experimental maximum velocity (with viscosity) calculated by interpolation.

The 3^rd^ part is related to the myosin head and its geometry: δX_Max_, δX_T_, δX_pre_ are linear ranges defining zones and domains associated with the angular ranges δθ_Max_, δθ_T_, δθ_pre_ corresponding to lever rotations.

The 4^th^ part refers to the time parameters of the cross-bridge cycle reactions: τ_preSB_ and τ_SB_ are the occurrence delay and time constant of the {SB} event, i.e. strong binding; τ_startF_, τ_preS_, τ_startS_, τ_preVS_, τ_preVS_, τ_startVS_ are the occurrence delays and time constants of the 3 events {startF}, {startS} and {startVS}, i.e. WS fast, slow and very slow initiations; τ_preSDE_ and τ_SDE_ are the occurrence delay and time constant of the {SlowDE} event, i.e. slow detachment.

r^2^ is the determination coefficient defined in the Methods section.

### Influence of the initial length of sarcomere (Fig 3 and Table 1)

During the isometric tetanus plateau preceding the force step, the fiber is in isometric conditions, i.e. at a fixed length (L0) which determines the average length of each sarcomere (L0_s_) such that:

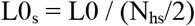

The Force/Velocity relationship is tested for 4 L0_s_ values equal to 1.85, 2.1, 2.3 and 2.6 μm.

If we compare the 2 columns “REF” and “2.1 μm” of Table 1 where L0_s_ = 2.1 μm in both cases, the figures are similar except for the values related to the fiber itself.

The data concerning the geometry of the myosin head (δX_Max_, δX_T_, δX_pre_) are identical for all 4 lengths, except the value of δX_T_ equal to 7 nm for L0_s_=1.85 μm, a decrease explained by the overlap of the myosin and actin filaments which limits the potential number of cross-bridges [9].

The time data relating to the chemical reactions of the cross-bridge cycle are identical except for the value of τ_SB_: the lowest value is found for L0_s_=2.1 μm, indicating an optimal realisation of the event {SB} or strong binding compared to the other 3 lengths; this efficiency criterion is to be associated with the maximum value of T0.

As τ_preSB_ is equal to 2.5 ms in all 4 cases, the theoretical maximum speeds are identical according to (16a) and (16b). The result is in accordance with the hypothesis formulated in Paper 4 that the massive elements (Z-disks associated with actin filaments and M-disks associated with myosin filaments) are the structures responsible for viscosity actions; since WS myosin heads play a negligible role, their number can be variable. Also the same value of 0.01 ms·nm^-1^ is assigned to ε for all 4 lengths (Table 1). After interpolation, the maximum observed velocities are close for the 4 plots (Fig 3 and Table 1). A difference of more than 12% between u_Max_ and u_Max,th_ is found, indicating an influence of viscosity if u > 1 nm·ms^-1^ per hs or if V > 1 L·s^-1^ for the fiber.

**Fig 3.**
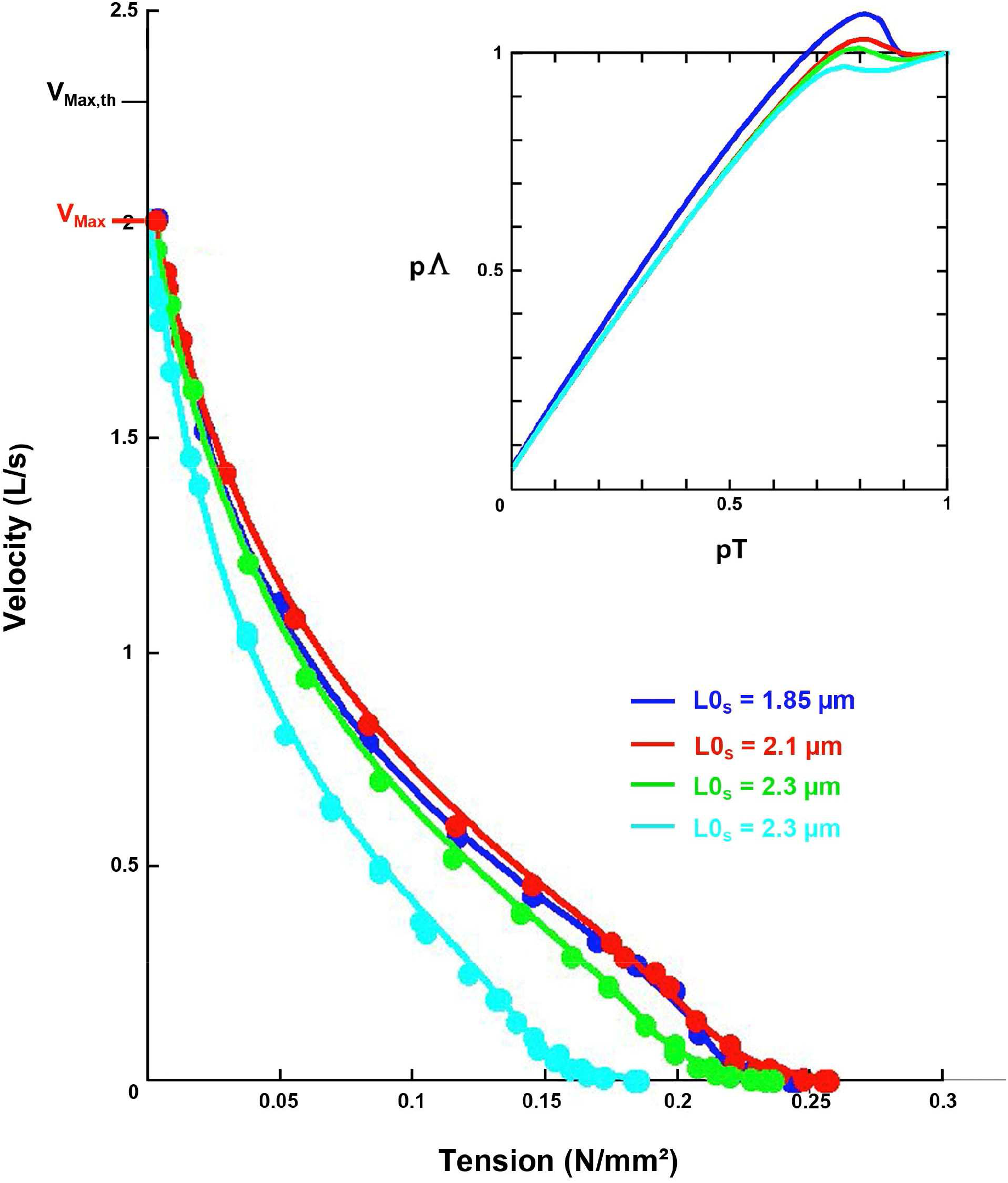
Force-Velocity relationships for 4 initial lengths of sarcomere. The 4 curves are drawn with a continuous line according to equation (12). L is a unit referring to the initial length of the fiber under standard conditions, i.e. V measured in mm/s is divided by “L0 = 8.35 mm” in all 4 cases. The points come from Figs 3A, 3B, 3C and 3D in [8]. Inset: the relationship of the relative number of WS heads (pΛ) as a function of the relative tension (pT) for the 4 lengths.

In the inset of Fig 3, at tensions close to 0.8·T0, the peak of the number of WS heads decreases in particular according to the p_startF_ value. The 3 plots of pΛ/pT relating to 2.1, 2.3 and 2.6 μm are similar to that of the reference graph in Fig 2. The plot relating to L0_s_ = 1.85 μm is above the other 3, especially for pT < 70% where the difference is due to the decrease in the range δX_T_.

### Influence of the inter-filament distance (Fig 4 and Table 2)

The increase in the tonicity of the *Ringer* solution from 1*R* to 1.44*R* osmotically compresses the cell fluid and reduces the inter-filament space.

The values in column “1*R*” (Table 2) are those of a standard *Ringer* solution administered to the fiber; they are close to those of the reference graph; see column “REF” (Table 1).

**Table 2.**
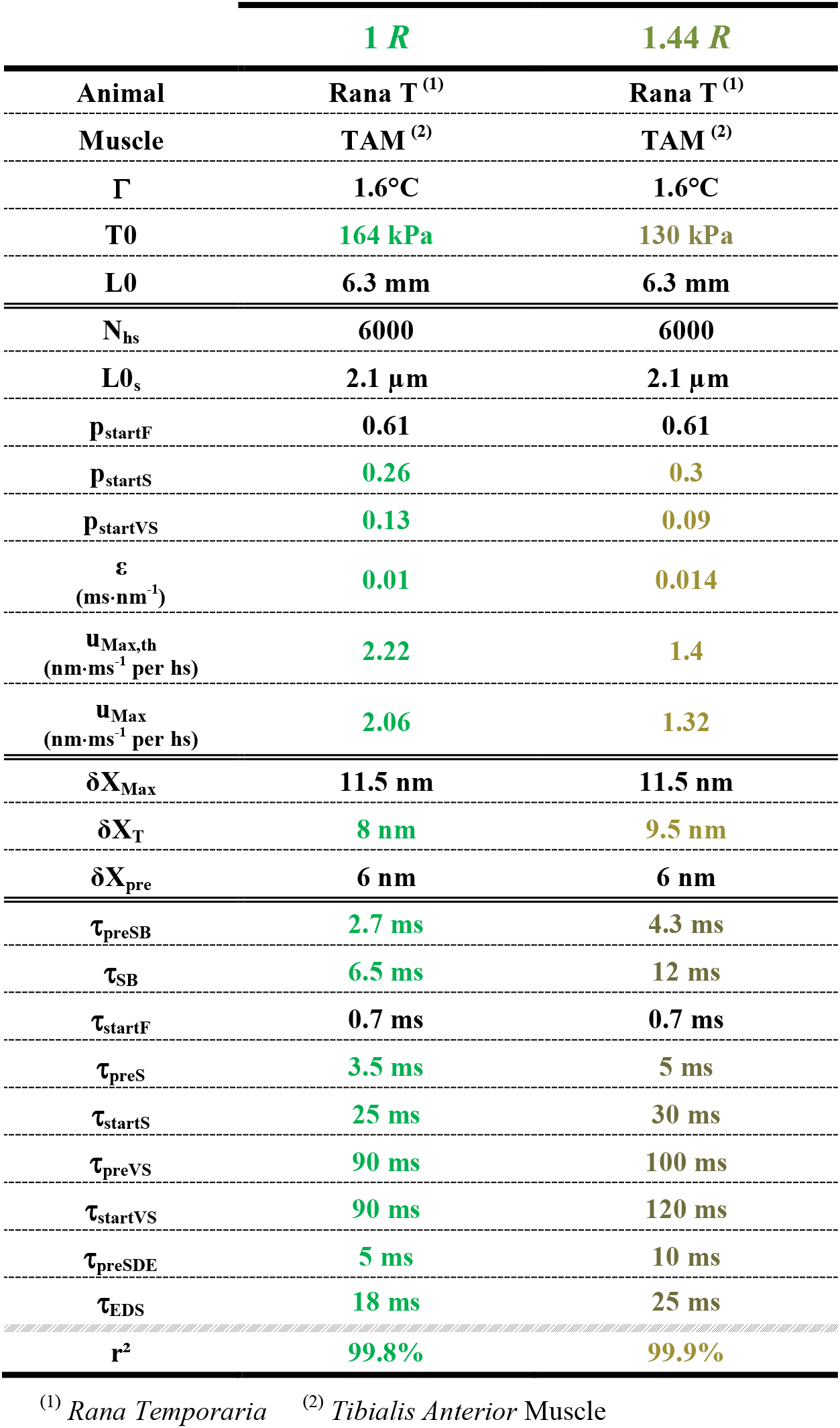
Parameter values for 2 tonicities of the *Ringer* solution in relation to the plots in Fig 4.

The hypertonicity of the *Ringer* solution (1.44 *R*) is analyzed in sub-paragraph J.16.7 of Supplement S4.J of Paper 4 with respect to a fiber isolated from the *tibialis anterior* of *rana Esculenta* [10]. The decrease in the inter-filament distance implies an increase of the angular range δθ_T_ and the viscous factor ε. The two species are close, so we use the figures found:

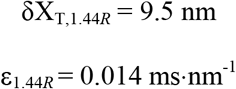

In column 1.44*R* (Table 3), the time parameters, τ_preSB_ and τ_SB_, τ_preS_ and τ_startS_, τ_preVS_, τ_preSDE_ and τ_SDE_, are higher than those for 1*R*. The shortening of the distance between the adjacent actin and myosin filaments reduces the number of strong binding and WS opportunities: this result in a 20% and 35% decrease in T0 and V_Max_ (Fig 4).

**Fig 4.**
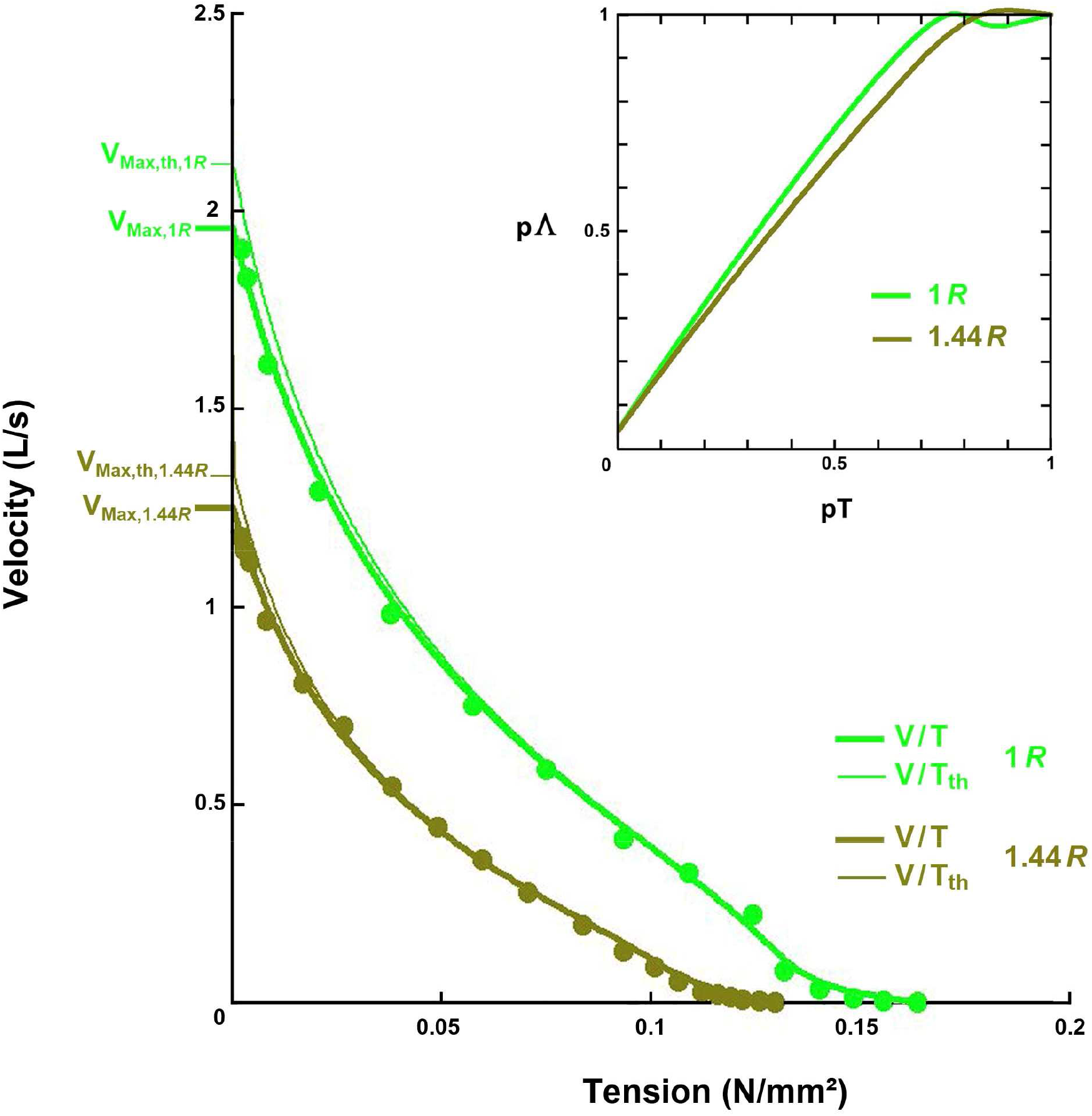
Force-Velocity relationships for 2 tonicities of the *Ringer* solution. The thick and thin line curves correspond to the V/T and V/T_th_ relationships according to equations (12) and (6), respectively. with a light green line for a standard *Ringer* solution (1*R*) and a dark green line for a hypertonic solution (1.44*R*). The points are from Figs 4A and 4B in [8]. Inset: the relation of the relative number of heads in WS (pΛ) as a function of the relative tension (pT).

**Table 3.**
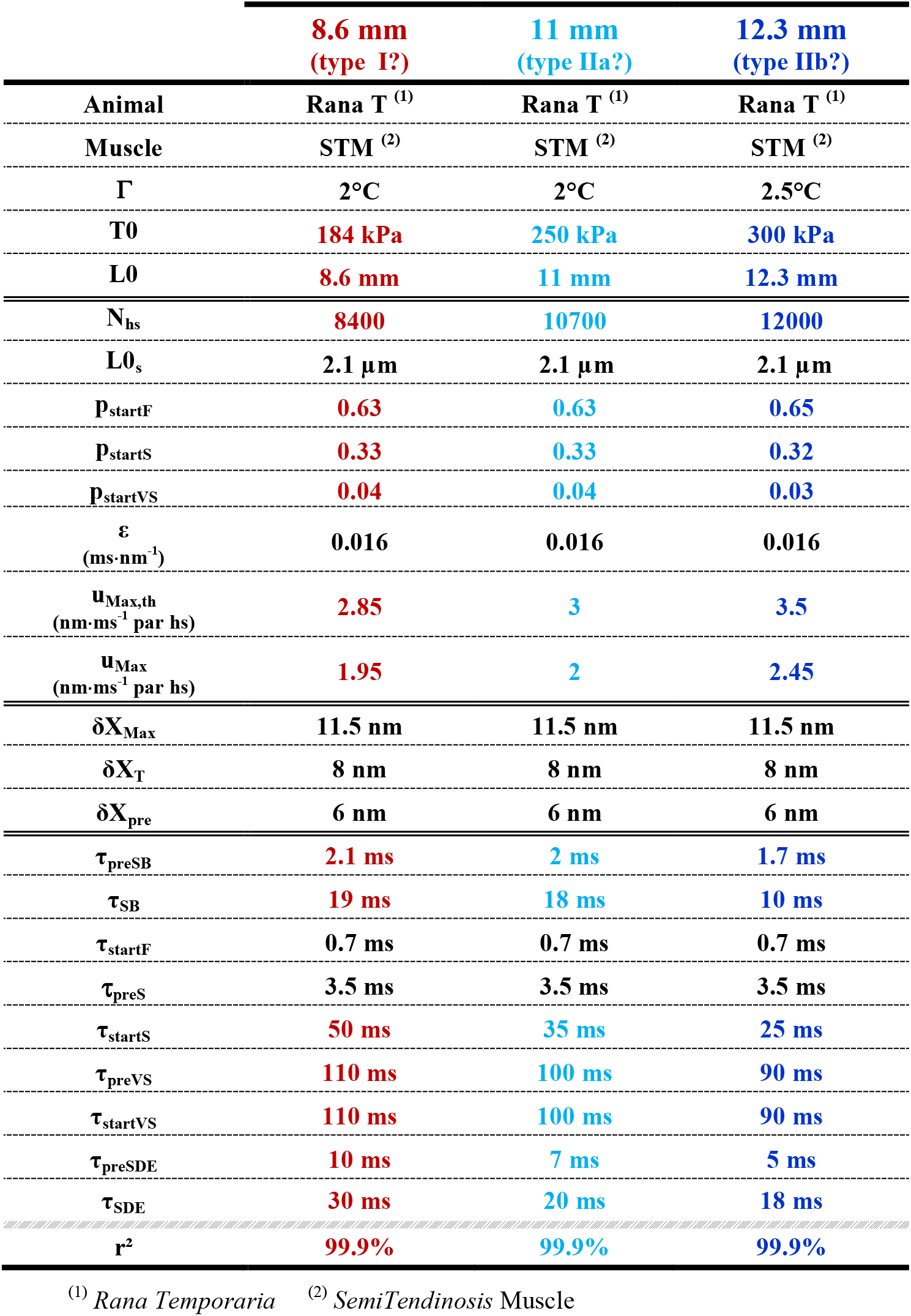
Parameter values for 3 fiber lengths in relation to the plots in Fig 5.

|  | 8.6 mm<br>(type I?) | 11 mm<br>(type IIa?) | 12.3 mm<br>(type IIb?) |
| --- | --- | --- | --- |
| Animal | Rana T <sup>(1)</sup> | Rana T <sup>(1)</sup> | Rana T <sup>(1)</sup> |
| Muscle | STM <sup>(2)</sup> | STM <sup>(2)</sup> | STM <sup>(2)</sup> |
| $\Gamma$ | 2°C | 2°C | 2.5°C |
| T0 | 184 kPa | 250 kPa | 300 kPa |
| L0 | 8.6 mm | 11 mm | 12.3 mm |
| $N_{hs}$ | 8400 | 10700 | 12000 |
| $L0_s$ | 2.1 $\mu\text{m}$ | 2.1 $\mu\text{m}$ | 2.1 $\mu\text{m}$ |
| $P_{startF}$ | 0.63 | 0.63 | 0.65 |
| $P_{startS}$ | 0.33 | 0.33 | 0.32 |
| $P_{startVS}$ | 0.04 | 0.04 | 0.03 |
| $\epsilon$<br>($\text{ms}\cdot\text{nm}^{-1}$) | 0.016 | 0.016 | 0.016 |
| $u_{Max,th}$<br>($\text{nm}\cdot\text{ms}^{-1}$ par hs) | 2.85 | 3 | 3.5 |
| $u_{Max}$<br>($\text{nm}\cdot\text{ms}^{-1}$ par hs) | 1.95 | 2 | 2.45 |
| $\delta X_{Max}$ | 11.5 nm | 11.5 nm | 11.5 nm |
| $\delta X_T$ | 8 nm | 8 nm | 8 nm |
| $\delta X_{pre}$ | 6 nm | 6 nm | 6 nm |
| $\tau_{preSB}$ | 2.1 ms | 2 ms | 1.7 ms |
| $\tau_{SB}$ | 19 ms | 18 ms | 10 ms |
| $\tau_{startF}$ | 0.7 ms | 0.7 ms | 0.7 ms |
| $\tau_{preS}$ | 3.5 ms | 3.5 ms | 3.5 ms |
| $\tau_{startS}$ | 50 ms | 35 ms | 25 ms |
| $\tau_{preVS}$ | 110 ms | 100 ms | 90 ms |
| $\tau_{startVS}$ | 110 ms | 100 ms | 90 ms |
| $\tau_{preSDE}$ | 10 ms | 7 ms | 5 ms |
| $\tau_{SDE}$ | 30 ms | 20 ms | 18 ms |
| $r^2$ | 99.9% | 99.9% | 99.9% |

<sup>(1)</sup> *Rana Temporaria* <sup>(2)</sup> *SemiTendinosis* Muscle

The narrowing of the inter-filament space increases the flow resistance. However, since the maximum speed is lower, the influence of viscosity remains minimal for 1.44*R*.

In the inset of Fig 4, for pT < 80%, the pΛ/pT plot for 1.44*R* is below that for 1*R*; the explanation comes from the increase in δX_T_.

### Influence of the fiber length or number of sarcomeres (Fig 5 and Table 3)

In sub-paragraph J.16.8 of Supplement S4.J to Paper 4, a study concerning a fiber isolated from the *semi-tendinosus* muscle of *Rana Temporaria* is conducted from Fig 6B in [11]. The value of the viscous coefficient is provided in conclusion:

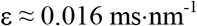

As the study in this paragraph refers to the points from Fig 3 in [5].where the three fibers tested belong to the *semi-tendinosus* of *R. Temporaria,* we assume that this ε value is common to them (Table 3). The V/T and V/T_th_ relationships for 3 initial fiber lengths (L0) are shown in Fig 5. We note that the difference between V_Max_ and V_Max,th_ enhances with L0; indeed, the τ_preSB_ delay decreases (Table 3) with the consequence that u_Max,th_ raises according to (16a), an increase that accentuates the role of viscosity via equation (11).

**Fig 5.**
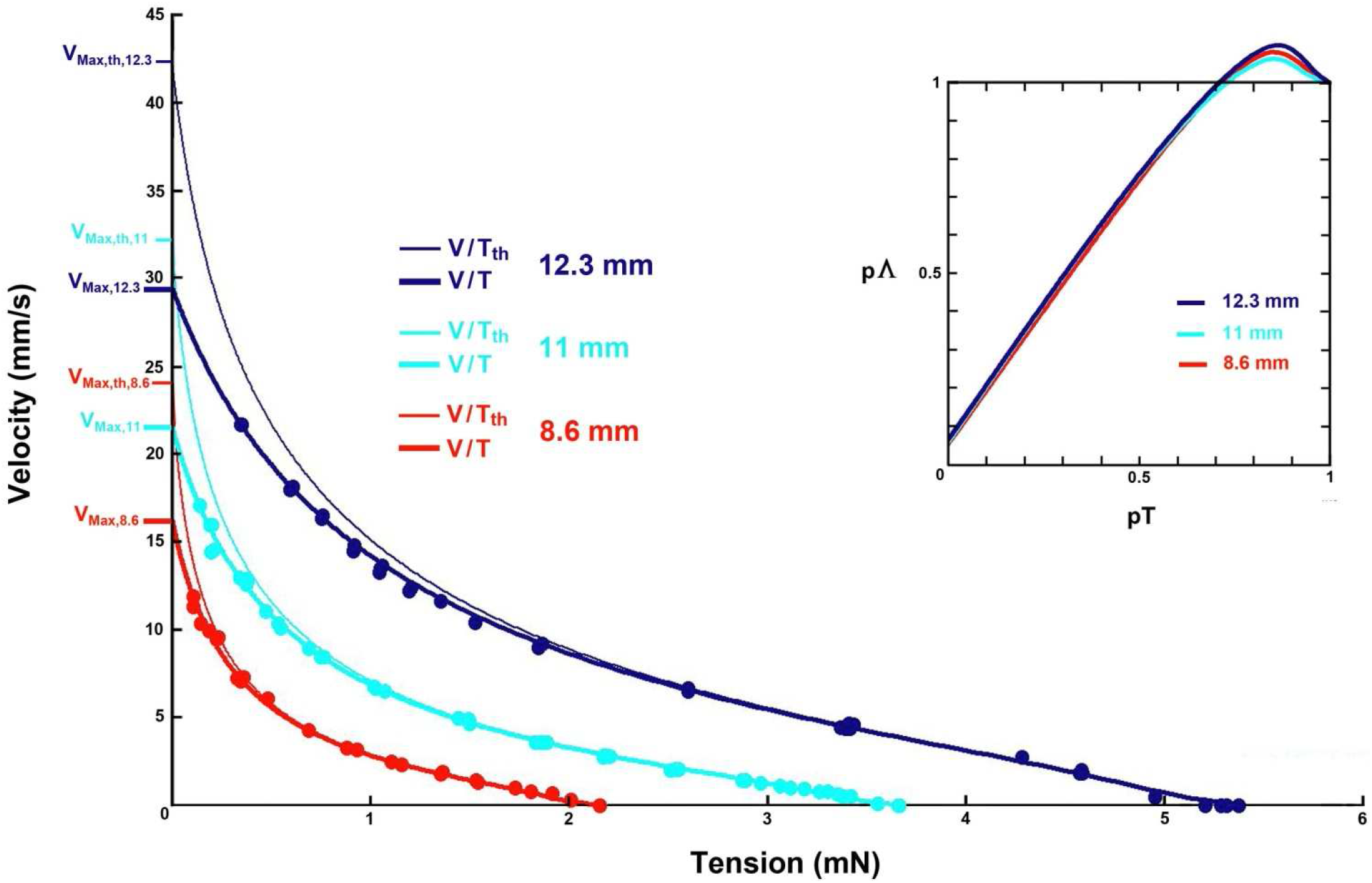
Force-Velocity relationships for 3 fiber lengths. The thick and thin line curves correspond to the V/T and V/T_th_ relationships according to equations (12) and (6), respectively. The points are from Fig 3 in [5]. Inset: relations of the relative number of WS heads (pΛ) as a function of the relative tension (pT).

In the inset of Fig 4, at tensions close to (0.8·T0), the peak of the number of WS heads is pronounced in all three cases due to the high value of p_startF_. The 3 plots are superimposable and comparable to the one of the referent graph if pT < 70%.

### Influence of the typology?

According to mechanical and histochemical criteria corresponding to isoforms specific to the heavy and light chains of myosin, skeletal muscle fibers are schematically diversified into 3 types: slow fiber (type I), intermediate fiber (type IIa) and fast fiber (type IIb). A study carried out on the *semi-tendinosus* muscle of *R. Temporaria* indicates that the 3 types are present [12]. For the elaboration of the curves, the intrinsic geometric data of the myosin heads (δX_Max_, δX_T_, δX_pre_) are identical. On the other hand, the time parameters decrease with L0 (Table 3) indicating an increase in chemical kinetics within the cross-bridge cycle. With the increase of T0 as a function of L0, the mechanical and chemical parameters of Table 3 suggest a hypothetical classification of the 3 fibers according to the 3 types.

The conjectures relating to the importance of viscosity depending on the typology and the constancy of the coefficient ε are to be checked.

### Influence of the internal temperature of the fiber (Fig 6 and Table 4)

The time data in Table 4 for cross-bridge cycle chemical reactions are consistent: occurrence delays and constant times decrease, i.e. the rates of all reactions increase with temperature (Γ).

**Table 4.** Parameter values for 4 experimental temperatures in relation to the plots in Fig 6.

| $\Gamma$ | 2°C | 7°C | 14.5° C | 21° C |
| --- | --- | --- | --- | --- |
| Animal | Rana E <sup>(1)</sup> | Rana E <sup>(1)</sup> | Rana E <sup>(1)</sup> | Rana E <sup>(1)</sup> |
| Muscle | TAM <sup>(2)</sup> | TAM <sup>(2)</sup> | TAM <sup>(2)</sup> | TAM <sup>(2)</sup> |
| T0 | 155 kPa | 195 kPa | 230 kPa | 241 kPa |
| L0 | 5 mm | 6 mm | 6 mm | 6 mm |
| N<sub>hs</sub> | 4800 | 5700 | 5700 | 5700 |
| L0<sub>s</sub> | 2.1 µm | 2.11 µm | 2.11 µm | 2.11µm |
| p<sub>startF</sub> | 0.63 | 0.6 | 0.65 | 0.64 |
| p<sub>startS</sub> | 0.31 | 0.32 | 0.3 | 0.3 |
| p<sub>startVS</sub> | 0.06 | 0.08 | 0.05 | 0.06 |
| $\varepsilon$<br>(ms·nm<sup>-1</sup>) | 0.015 | 0.009 | 0.004 | 0.002 |
| u<sub>Max,th</sub><br>(nm·ms<sup>-1</sup> par hs) | 2.4 | 4.3 | 10 | 17.15 |
| u<sub>Max</sub><br>(nm·ms<sup>-1</sup> par hs) | 2.1 | 3.4 | 7.85 | 13.05 |
| δX<sub>Max</sub> | 11.5 nm | 11.5 nm | 11.5 nm | 11.5 nm |
| δX<sub>T</sub> | 8 nm | 8 nm | 8 nm | 8 nm |
| δX<sub>pre</sub> | 6 nm | 6 nm | 6 nm | 6 nm |
| τ<sub>preSB</sub> | 2.5 ms | 1.4 ms | 0.6 ms | 0.35 ms |
| τ<sub>SB</sub> | 6.5 ms | 4.5 ms | 1.6 ms | 1 ms |
| τ<sub>startF</sub> | 0.8 ms | 0.7 ms | 0.7 ms | 0.6 ms |
| τ<sub>preS</sub> | 3.5 ms | 3 | 2 ms | 1 |
| τ<sub>startS</sub> | 25 ms | 11 ms | 6 ms | 3.5ms |
| τ<sub>preVS</sub> | 80 ms | 60 ms | 40 ms | 30 ms |
| τ<sub>startVS</sub> | 90 ms | 70 ms | 40 ms | 30 ms |
| τ<sub>preSDE</sub> | 5 ms | 4 ms | 2.5 ms | 2 ms |
| τ<sub>SDE</sub> | 18 ms | 14 ms | 11 ms | 7 ms |
| r<sup>2</sup> | 99.8% | 99.8% | 99.8% | 99.9% |

<sup>(1)</sup> *Rana Temporaria* <sup>(2)</sup> *Tibialis Anterior* Muscle

For 2°C, the value of ε is equal to 0.015 ms·nm^-1^ (see Methods section). The values displayed in Table 4 for the other 3 temperatures are determined in sub-paragraph J.16.6 of Supplement S4.J of Paper 4.

As the temperature rises, the difference between V_Max_ and V_Max,th_ increases, affirming the importance of viscosity at high speeds even for low values of ε.

## Discussion

### Adequacy of the model

Experiments on muscle fiber have various biases. Tensions are sometimes expressed in Kilopascal (kPa), a unit of pressure equal to the ratio of a force to a surface; data in kPa should therefore be corrected by multiplying the pressure value by the cross-sectional area of the fiber, a geometric characteristic that is difficult to quantify and variable according to the tests. The rise to the isometric tetanus plateau prior to each force step can be different and influence the measurements, in particular by affecting the maximum proportions p_startF_, p_startS_ and p_startVS_. The repetition of the tests may modify the internal structure of the fiber and alter the results. The points are occasionally averages calculated from tests performed on several fibers, increasing the range of confidence intervals. Notwithstanding these error factors, a good agreement between experimental points and theoretical curves is found with r^2^ > 99.75% (Tables 1 to 4).

The adjustment chosen is not necessarily the best, other combinations of values offered higher determination coefficients. We have made our choices to be consistent with the usual data in the literature and those of accompanying papers. As a general rule, the values of δX_Max_, δX_T_ and δX_pre_ are equal to 11.5 nm, 8 nm and 6 nm, those of p_startF_ and p_startS_ are close to 60% and 30%, those of τ_startF_, τ_startS_, τ_startVS_ and τ_SDE_ are at low temperatures, respectively, equal to 0.7 ms, 25 ms, 90 ms and 18 ms (Table 1 to 4).

### Stroke size

The maximum angular variation (δθ_Max_) of the lever of a myosin II head between the *up* and *down* positions is typically measured as 70° [15,16,17,18,19,20,21,22,23]. With a lever length S1b slightly less than 10 nm [15,24], the relationship (1) makes it possible to determine the maximum linear range (δX_Max_):

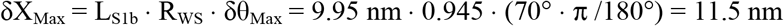

The acronym δX_Max_ represents the stroke size of a myosin II head whose usual value varies between 10 and 12 nm [17,25,26,27,28,29,30,31]. The value of δX_Max_ calculated above is standard in the tables of the article.

### Why does the stroke size seem shorter for high tensions?

Using the interference properties of X-rays, the angular dispersion (δθ_T_) of the levers of the myosin II heads in WS during the isometric tetanus plateau is analyzed and the measurement of δθ_T_ varies between 40° and 50° [31,32,33,34,35]. For the model, we adopted the value of 49° close to 50°.

The relationship (1) provides a linear displacement (δX_T_) corresponding to δθ_T_ equal to:

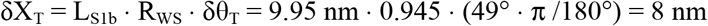

This value of δX_T_ is common to all the examples examined in the article with the exception of two special cases. M. Reconditi and his co-authors [35] indicate that the stroke size relatively to high tensions near T0 is estimated at 8 nm, and that this stroke size extends to a maximum of 13 nm for low tensions. H. Huxley and his co-authors [33] continue this research by presenting a maximum step equal to 7.5 nm for high tensions which increases to 11.2 nm for low tensions. Our model offers an explanation: the maximum step or stroke size (δX_Max_) is a constant but the {SlowDE} event which operates at very high tensions and very slow speeds allows the detachment of WS heads whose lever has an angular position between θ_down_ and θ_T_, i.e. an apparent stroke size for the heads remaining in WS corresponding to δX_T_. The more the tension decreases, the more the shortening speed increases, the less time the WS heads with a lever orientation between θ_down_ and θ_T_ have to detach, and the closer the apparent stroke size gets to δX_Max_, thus varying from 8 to 11.5 nm according to the data in our model.

### Average durations and delay times related to the cross-bridge cycle reactions

Temporal data relating, on the one hand, to the rapid, slow or very slow initiation of a WS, and on the other hand, to the slow detachment are explained in Supplement S1.B. Factors such as interfilament distance, typology and temperature interfere with reaction rates and associated time constants. The factorial changes in the theoretical values of these parameters are consistent with the experimental results.

### Biphasic aspect of the curve

The model provides a simple solution to the convexity marked for T > (0.78·T0): the {SlowDE} event occurs only at very slow and slow speeds (Figs 2, 3 and 4).

### Number of heads in WS per hs

The relationship of the relative number of WS heads per hs (pΛ) to the relative value of the force step (pT) is shown in the inserts in Figs 2 to 5. The behaviour of pΛ is variable for high tensions (pT>70%), the relative number decreases slowly from 1, sometimes it decreases then rises to 1 and sometimes it forms a pronounced peak above 1 if p_startF_>60%. For low and medium tensions (pT<70%), a quasi-linear decrease is observed up to low tensions.

Under the condition “p_startF_<60%”, the pΛ/pT plot is comparable to that in Fig 2B in [31] where the intensity of the M3 reflection of X-rays (IM3) is an indicator of the number of WS heads.

### The three factors limiting the maximum velocity of shortening

According to equality (16a), the theoretical maximum speed of a hs (uMax,th) is equal to the ratio of δX_pre_ to τ_preSB_, i.e. the extent necessary for the rapid initiation of a strong binding on the occurrence time of {SB}, the event that prefigures {startF}. Since δX_pre_ is a constant (see Tables 1 to 4), the first limiting factor is τ_preSB_.

In the expression (16b) applying to the theoretical maximum fiber velocity (V_Max,th_), the number of hs (N_hs_) is presented as a multiplier coefficient. Thus the length of the fiber is the second limiting factor, illustrated in Fig 5.

The theoretical maximum speeds u_Max,th_ and V_Max,th_ are higher than the maximum observed speeds u_Max_ and V_Max_ determined by interpolation of the curves u/pT and V/T. The cause is due to the presence of viscosity, presence quantified by the coefficient ε introduced in (11). The viscosity coefficient ε is the third limiting factor; see Figs 2, 4, 5 and 6.

**Fig 6.**
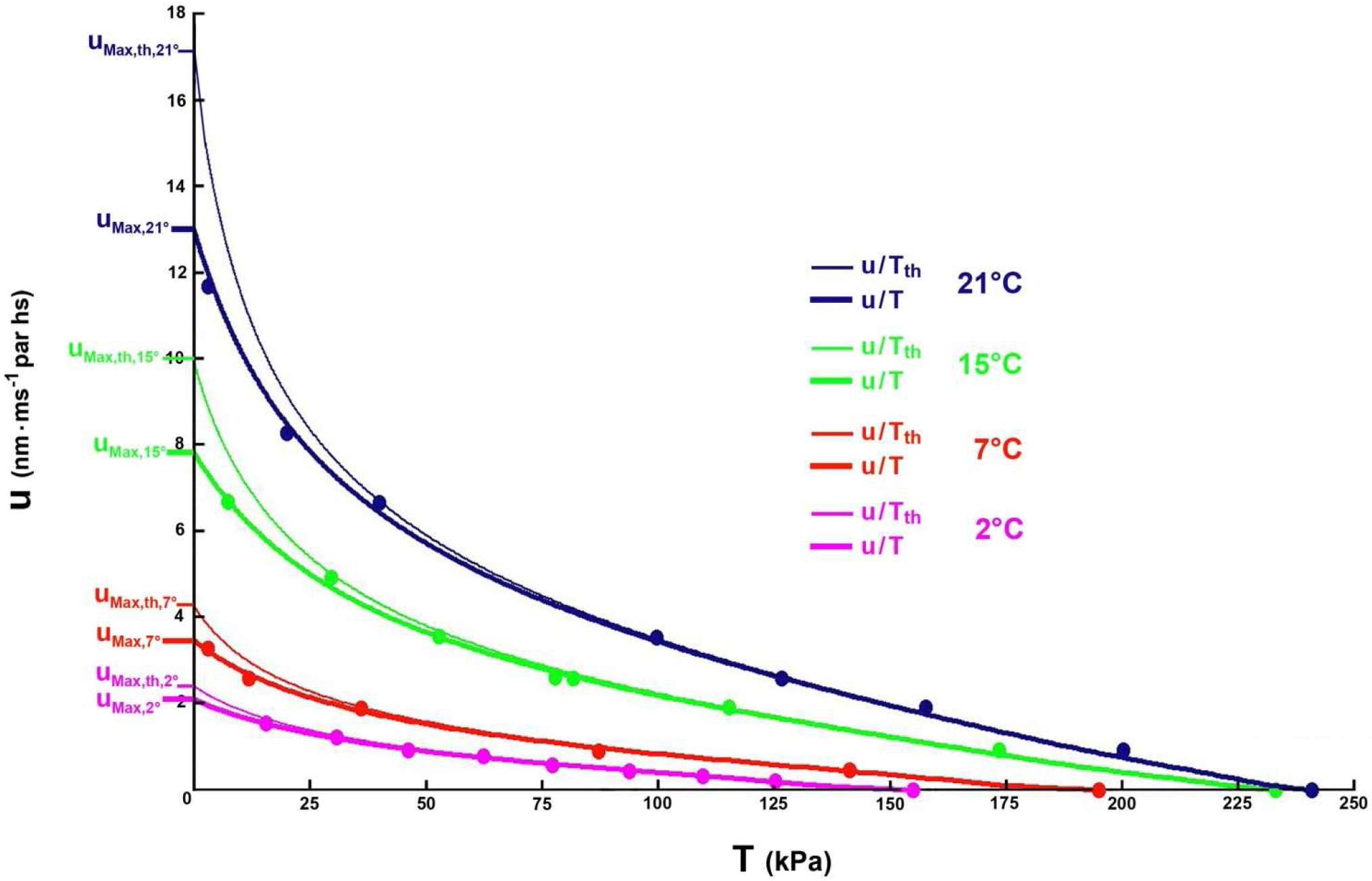
Force-velocity relationships for 4 experimental temperatures. The thick and thin line curves correspond to the V/T and V/T_th_ relationships according to equations (12) and (6), respectively. The fibers are isolated from the *tibialis anterior* muscle of *Rana Esculenta*. The purple dots are from Fig 3A in [13] and the red, green and blue dots from Fig 7A in [14].

### Presence of viscosity

The role of viscosity is particularly highlighted during phase 1 of a length step (see accompanying Paper 4) and impacts phase 4 of a force step when the fiber shortens at constant tension and velocity. For most physiologists, viscosity is absent from the Force/Velocity relationship.

A.V. Hill wrote p. 164 in [1]: “*It is difficult to think of a force of this kind (viscosity) which might be present in muscle*”.

L.E. Ford noted p. 133 in [36]: *“All the matches reported in this Paper were made with M_0_, representing viscosity in the fiber, set to zero*”.

R. Elongovan mentioned page 1237 in [14]: *“There is negligible viscous drag on the filaments as they are propelled by myosin along their axis*”.

Our model introduces the viscosity forces whose presence cannot be neglected at high speeds, especially for long fibers (Fig 5). Viscosity is shown to be a limiting factor in the maximum speed of shortening.

## Conclusion

A theoretical model of the Force/Velocity relationship is proposed in good adequacy with the experimental points identified in examples in the literature. The origin of the equations underlying the F-V relationship still needs to be clarified; their demonstration is one of the objectives of accompanying Papers 2 to 6.

## Supporting information

Supplementary Chapter. Probabilities and irreversible chemical reactions

Supplementary Chapter. Mathematization of the cross-bridge cycleComputer Programs for calculating and plotting Force/Velocity relationships

Computer Programs for calculating and plotting Force/Velocity relationships

Data used for computer programs of Paper 1

## Acknowledgements

I thank Professor K.A.P. Edman for allowing me to reproduce the points presented in Figs 2 to 5.

I thank Professor V. Lombardi for allowing me to reproduce the points presented in Fig 6.

## Supporting information

**S1.A Supplementary Chapter. Probabilities and irreversible chemical reactions.**

A.1 Random variable

A.2 Indicator functions

A.3 Mathematical reminders

A.4 Density of the sum of 2 independent random variables each following an exponential law

A.5 Density of the sum of 3 independent random variables each following an exponential law

A.6 Counting

A.7 Succession of n irreversible chemical reactions

A.8 Succession of n irreversible chemical reactions where the number of steps is unknown

A. 9 Conclusion

References of Supplement S1.A

**S1.B Supplementary Chapter. Mathematization of the cross-bridge cycle.**

B.1 The myosin head as a mechanical object

B.2 The myosin head as a chemical reactor

B.3 Initiation of the working stroke (WS)

B.4 Preparation of the {startF} event by the {SB} event

B.5 Realisation of the Detachment state according to the WS state

B.6 Fast Detachment

B.7 Slow Detachment

References of Supplement S1.B

**CP1 Supplementary Material. Computer Programs for calculating and plotting Force/Velocity relationships.** Algorithms are written in Visual Basic 6 © Microsoft.

**DA1 Supplementary Material. Data displayed in the tables to Paper 1 and used for computer programs of Supplement CP1.** Access Tables are transferred to Excel sheets.

