## Supplementary Chapter. Probabilities and irreversible chemical reactions for "Mechanical model of muscle contraction. 1. Force-velocity relationship"

### S1.A Supplementary Chapter of Paper 1

### Probabilities and irreversible chemical reactions

##### A.1 Random variable

We introduce ω as an elementary event of an experience and Ω as the set of all the elementary events of this experience. For each elementary event ω, a real x is matched. The random variable X is defined as the bijective application of Ω in R such that:

$$x = X(\omega) \tag{A1}$$

with $\omega \in \Omega$ and $x \in R$.

Similarly, the random variable T is associated with time t and the random variable Θ with the angle θ. In theory, the real x, t and θ are dimensionless but in practice they present a unit of measurement such as the millisecond for t or the radian for θ. To overcome this disadvantage, it is sufficient to divide the real measured by its unit of measurement.

#### A.2 Indicator functions

We submit K as a subset of elementary events from Ω. We note $\mathbb{1}_K$ the indicator function of the elementary event ω related to K as:

$$\mathbb{1}_K(\omega) = \begin{cases} 1 & \text{si } \omega \in K \\ 0 & \text{unless} \end{cases} \tag{A2a}$$

We present S as a subset of R corresponding to the subset K in Ω according to random variable X defined in (A1). We note $\mathbb{1}_S$ the indicator function of the real x relative to S such that:

$$\mathbb{1}_S(x) = \begin{cases} 1 & \text{si } x \in S \\ 0 & \text{unless} \end{cases} \tag{A2b}$$

#### A.3 Mathematical reminders

We introduce x as a real and $a_1$, $a_2$ and $a_3$ as non-zero positive constants with $a_1 \neq a_2$, $a_1 \neq a_3$ and $a_2 \neq a_3$.

The Taylor series for the exponential function is:

$$e^x = 1 + \frac{x}{1!} + \frac{x^2}{2!} + \frac{x^3}{3!} + ... + \frac{x^n}{n!} + ... \quad \text{(A3)}$$

**Let the function $p_1(x)$:**

$$p_1(x) = 1 - e^{-\frac{x}{a_1}} \quad \text{(A4a)}$$

Function $p_1$ is expanded to the 1st order. If $x \to 0$, we check:

$$p_1(x) \to \frac{x}{a_1} \quad \text{(A4b)}$$

**Let the function $p_2(x)$:**

$$p_2(x) = 1 - \frac{a_1 \cdot e^{-\frac{x}{a_1}}}{(a_1 - a_2)} - \frac{a_2 \cdot e^{-\frac{x}{a_2}}}{(a_2 - a_1)} \quad \text{(A5a)}$$

Function $p_2$ is expanded to the 2nd order. If $x \to 0$, we check:

$$p_2(x) \to \frac{x^2}{2 \cdot a_1 \cdot a_2} \quad \text{(A5b)}$$

**Let the function $p_3(x)$:**

$$p_3(x) = 1 - \frac{a_1^2 \cdot e^{-\frac{x}{a_1}}}{(a_1 - a_2) \cdot (a_1 - a_3)} - \frac{a_2^2 \cdot e^{-\frac{x}{a_2}}}{(a_2 - a_1) \cdot (a_2 - a_3)} - \frac{a_3^2 \cdot e^{-\frac{x}{a_3}}}{(a_3 - a_2) \cdot (a_3 - a_1)} \quad \text{(A6a)}$$

The $p_3$ function is expanded to the 3rd order. If $x \to 0$, we check:

$$p_3(x) \to \frac{x^3}{6 \cdot a_1 \cdot a_2 \cdot a_3} \quad \text{(A6b)}$$

The functions $p_4(x)$, $p_5(x)$, ... , $p_n(x)$ are constructed according to a recurrent pattern. Examples illustrating the behaviour of functions $p_1$, $p_2$, $p_3$, $p_4$ and $p_5$ are shown in Figure A1a for specific values of $a_1$, $a_2$, $a_3$, $a_4$ and $a_5$.

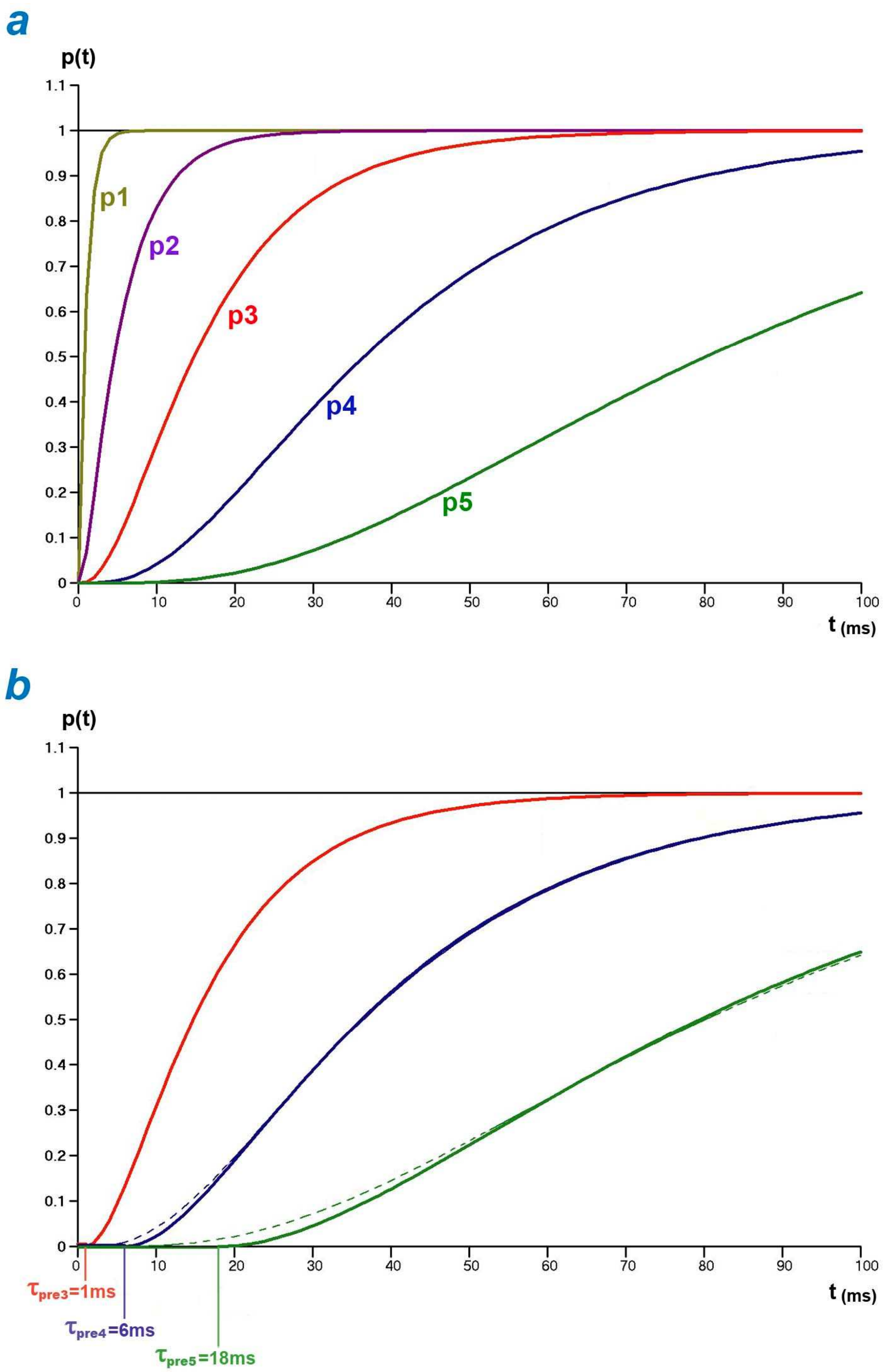


**Fig A1. Example of functions p1, p2, p3, p4 and p5.**

The 5 coefficients, $a_1$, $a_2$, $a_3$, $a_4$ and a5, are equal to 1 ms, 5 ms, 12 ms, 25 ms and 50 ms, respectively.(a) Temporal evolutions of the 5 functions. (b) Approximations of $p_3$, $p_4$ and $p_5$ according to (A21) with $\tau_{pre3}= a_1$, $\tau_{pre4}= (a_1+a_2)$ and $\tau_{pre5}= (a_1+a_2+a_3)$, respectively. The $p_3$, $p_4$ and $p_5$ plots appear with a dotted line.

#### A.4 Density of the sum of 2 independent random variables following each an exponential law

We present $T_1$ and $T_2$ as 2 random variables associated with the real t each following an exponential law of respective parameters $\lambda_1$ and $\lambda_2$ with $\lambda_1 \neq \lambda_2$. The probability densities of $T_1$ and $T_2$, noted respectively $f_1$ and $f_2$, are:

$$f_1(t) = \lambda_1 \cdot e^{-\lambda_1 \cdot t} \cdot \mathbf{1}_{R_+}(t) \qquad \text{(A7a)}$$

$$f_2(t) = \lambda_2 \cdot e^{-\lambda_2 \cdot t} \cdot \mathbf{1}_{R_+}(t) \qquad \text{(A7b)}$$

where t is a positive real representing the instantaneous time; $\mathbf{1}$ is the indicator function defined in (A2b).

If $T_1$ and $T_2$ are 2 independent random variables, the probability density of the sum of $T_1$ and $T_2$ ($f_{1+2}$) is equal to the convolution product of $f_1$ and $f_2$ [1]:

$$f_{1+2}(t) = (f_1 * f_2)(t) = \int_{-\infty}^{+\infty} f_1(t-z) \cdot f_2(z) \cdot dz$$

where " * " is the symbol for the convolution product.

The previous expression is reformulated with (A7a) and (A7b):

$$f_{1+2}(t) = \int_{-\infty}^{+\infty} \left[\lambda_1 \cdot e^{-\lambda_1 \cdot (t-z)} \cdot \mathbf{1}_{R_+}(t-z) \cdot \lambda_2 \cdot e^{-\lambda_2 \cdot z} \cdot \mathbf{1}_{R_+}(z)\right] \cdot dz \qquad \text{(A8)}$$

The product of the 2 indicator functions is rewritten [2]:

$$\mathit{1}_{R_+}(t-z) \cdot \mathit{1}_{R_+}(z) = \begin{cases} 1 & \text{if} \quad z \geq 0 \quad \text{and} \quad (t-z) \geq 0 \\ 0 & \text{unless} \end{cases}$$

$$= \begin{cases} \mathbf{1}_{[0;t]}(z) & \text{if} \quad t \geq 0 \\ 0 & \text{unless} \end{cases}$$

$$= \mathbf{1}_{R_+}(t) \cdot \mathbf{1}_{[0;t]}(z) \qquad \text{(A9)}$$

By entering (A9) in (A8), we obtain:

$$f_{1+2}(t) = \mathbf{1}_{R_+}(t) \cdot \left(\lambda_1 \cdot \lambda_2 \cdot e^{-\lambda_1 \cdot t}\right) \cdot \left(\int_{-\infty}^{+\infty} e^{-z \cdot (\lambda_2 - \lambda_1)} \cdot \mathbf{1}_{[0;t]}(z) \cdot dz\right)$$

With the definition of an indicator function:

$$f_{1+2}(t) = \mathbf{1}_{R_+}(t) \cdot \left(\lambda_1 \cdot \lambda_2 \cdot e^{-\lambda_1 \cdot t}\right) \cdot \left(\int_0^t e^{-z \cdot (\lambda_2 - \lambda_1)} \cdot dz\right)$$

And after integration:

$$f_{1+2}(t) = \left(\frac{\lambda_1 \cdot \lambda_2 \cdot e^{-\lambda_1 \cdot t}}{\lambda_2 - \lambda_1}\right) \cdot \left[1 - e^{-t \cdot (\lambda_2 - \lambda_1)}\right]$$

The probability density of the sum of 2 random variables following each an exponential law is formulated:

$$f_{1+2}(t) = \frac{\lambda_1 \cdot \lambda_2}{(\lambda_2 - \lambda_1)} \cdot e^{-\lambda_1 \cdot t} + \frac{\lambda_1 \cdot \lambda_2}{(\lambda_1 - \lambda_2)} \cdot e^{-\lambda_2 \cdot t} \qquad \text{(A10)}$$

This result is obtained directly with the product of the characteristic functions of densities $f_1$ and $f_2$ (see the next paragraph with case n=3). This demonstration was chosen because the rewriting of the indicator product in (A9) underlines that the occurrence of one of the 2 events precedes that of the other and applies well to two successive events in time.

##### A.5 Density of the sum of 3 independent random variables following each an exponential law

The characteristic function of a random variable is defined by the Fourier transform of the density of this random variable:

$$\mathcal{F}[f(x)] = \int f(t) \cdot e^{i \cdot x \cdot t} \cdot dt \qquad \text{(A11)}$$

where i represents the unit of pure imaginary numbers.

Let be $T_1$, $T_2$ and $T_3$, 3 random variables each following an exponential law of parameters respectively $\lambda_1$, $\lambda_2$ and $\lambda_3$ with $\lambda_1 \neq \lambda_2$ , $\lambda_1 \neq \lambda_3$ and $\lambda_2 \neq \lambda_3$.

The probability densities of $T_1$, $T_2$ and $T_3$ are written:

$$f_1(t) = \lambda_1 \cdot e^{-\lambda_1 \cdot t} \cdot \mathbf{1}_{R_+}(t) \qquad \text{(A12a)}$$

$$f_2(t) = \lambda_2 \cdot e^{-\lambda_2 \cdot t} \cdot \mathbf{1}_{R_+}(t) \qquad \text{(A12b)}$$

$$f_3(t) = \lambda_3 \cdot e^{-\lambda_3 \cdot t} \cdot \mathbf{1}_{R_+}(t) \qquad \text{(A12c)}$$

We consider the random variable T sum of $T_1$, $T_2$ and $T_3$ and the density of T ($f_{1+2+3}$). The characteristic function of the density of the sum of 3 independent random variables is equal to the product of the characteristic functions of the 3 densities of these 3 random variables [1,3].

From the above, with (A11), (A12a), (A12a), (A12b) and (A12c) and by distributivity of the sum and the convolution product, the characteristic function of $\mathbf{f_{1+2+3}}$ is formulated:

$$\mathcal{F}[f_{1+2+3}(x)] = \left( \int_0^\infty \lambda_1 \cdot e^{-(\lambda_1 - i \cdot x) \cdot t} \cdot dt \right) \cdot \left( \int_0^\infty \lambda_2 \cdot e^{-(\lambda_2 - i \cdot x) \cdot t} \cdot dt \right) \cdot \left( \int_0^\infty \lambda_3 \cdot e^{-(\lambda_3 - i \cdot x) \cdot t} \cdot dt \right)$$

Integration gives:

$$\mathcal{F}[f_{1+2+3}(x)] = \frac{\lambda_1}{(\lambda_1 - i \cdot x)} \cdot \frac{\lambda_2}{(\lambda_2 - i \cdot x)} \cdot \frac{\lambda_3}{(\lambda_3 - i \cdot x)}$$

After development:

$$\mathcal{F}[f_{1+2+3}(x)]=\frac{\lambda_1\cdot\lambda_2\cdot\lambda_3}{(\lambda_2-\lambda_1)\cdot(\lambda_3-\lambda_1)}\cdot\left(\frac{1}{\lambda_1-i\cdot x}\right)+\frac{\lambda_1\cdot\lambda_2\cdot\lambda_3}{(\lambda_1-\lambda_2)\cdot(\lambda_3-\lambda_2)}\cdot\left(\frac{1}{\lambda_2-i\cdot x}\right)+\frac{\lambda_1\cdot\lambda_2\cdot\lambda_3}{(\lambda_1-\lambda_3)\cdot(\lambda_2-\lambda_3)}\cdot\left(\frac{1}{\lambda_3-i\cdot x}\right)$$

By going through the inverse transformation of the Fourier transform, we obtain the density of the sum of 3 independent random variables:

$$f_{1+2+3}(t)=\frac{\lambda_1\cdot\lambda_2\cdot\lambda_3}{(\lambda_2-\lambda_1)\cdot(\lambda_3-\lambda_1)}\cdot e^{-\lambda_1\cdot t}+\frac{\lambda_1\cdot\lambda_2\cdot\lambda_3}{(\lambda_1-\lambda_2)\cdot(\lambda_3-\lambda_2)}\cdot e^{-\lambda_2\cdot t}+\frac{\lambda_1\cdot\lambda_2\cdot\lambda_3}{(\lambda_1-\lambda_3)\cdot(\lambda_2-\lambda_3)}\cdot e^{-\lambda_3\cdot t} \qquad \text{(A13)}$$

#### A. 6 Counting

Let N myosin heads concerned by a K event whose probability of occurrence between the times 0 and t is noted p. Let us consider N independent random variables, generically named $A_b$ (with b = 1, 2, ... , N) which correspond to the occurrence time of K between 0 and t for each of the N heads.

Let $B_b$ the Bernouilli's random variable equal to $\mathbb{1}_K(t)$ for the head n° b. The N $B_b$ (with b = 1, 2,..., N) are independent Bernouilli's random variables of parameter p. By definition the sum of the N $B_b$ is a random variable named $N_K$ which follows a binomial law of parameters N and p. It is deduced that the average number of myosin heads for which K has occurred between 0 and t is equal to:

$$E(N_K) = N\cdot p \qquad \text{(A14)}$$

where E is the expectation of $N_K$.

**Special case 1**

Let K, an event occurring at random in time; the probability density of K is an exponential law of parameter λ with $f(t)=\lambda\cdot e^{-\lambda\cdot t}$. The N random variables $A_b$ (with b = 1, 2, ... , N), all concerned by K event, follow independently the same density f(t). The probability p of K occurrence before t for the head n° b is by definition equal to:

$$p=\int_0^t \lambda\cdot e^{-\lambda\cdot z}\cdot dz=1-e^{-\lambda\cdot t}$$

By applying (A14), we obtain:

$$E_1(N_K)=N\cdot\left(1-e^{-\lambda\cdot t}\right) \qquad \text{(A15)}$$

We recognize $p_1(t)/N$ where the function $p_1$ has been introduced in (A4a) with $\lambda_1=1/a_1$.

**Special case 2**

Let K, the succession of 2 events $K_1$ and $K_2$ each occurring at random whose respective probability densities are $f_1$ and $f_2$, the 2 exponential functions defined in (A7a) and (A7b) ; the probability density of K is $f_{1+2}$ determined in (A10). The N random variables $A_b$ follow independently the same density $f_{1+2}$. The probability p of K occurrence before t for the head n° b is by definition:

$$p = \int_0^t f_{1+2}(z) \cdot dz$$

After the integration provided in (A10) and with (A14):

$$E_2(N_K) = N \cdot \left[ 1 - \frac{\lambda_2}{\lambda_2 - \lambda_1} \cdot e^{-\lambda_1 \cdot t} - \frac{\lambda_1}{\lambda_1 - \lambda_2} \cdot e^{-\lambda_2 \cdot t} \right] \qquad \text{(A16)}$$

We recognize $p_2(t)/N$ where the function $p_2$ has been introduced in (A5a) with $\lambda_1=1/a_1$ and $\lambda_2=1/a_2$.

**Special case 3**

Let K, the succession of 3 events $K_1$, $K_2$ and $K_3$, each occurring at random, whose respective probability densities are $f_1$, $f_2$ and $f_3$, the 3 exponential functions defined in (A12a), (A12b) and (A12c); the probability density of K is $f_{1+2+3}$ determined in (A13). The N random variables $A_b$ follow independently the same probability density law $f_{1+2+3}$. By definition, the probability p of K occurrence before t for the head n° b is formulated according to:

$$p = \int_0^t f_{1+2+3}(z) \cdot dz$$

After the integration delivered in (A13) and with (A14):

$$E_3(N_K) = N \cdot \left[ 1 - \frac{\lambda_2 \cdot \lambda_3 \cdot e^{-\lambda_1 \cdot t}}{(\lambda_2 - \lambda_1) \cdot (\lambda_3 - \lambda_1)} - \frac{\lambda_1 \cdot \lambda_3 \cdot e^{-\lambda_2 \cdot t}}{(\lambda_1 - \lambda_2) \cdot (\lambda_3 - \lambda_2)} - \frac{\lambda_1 \cdot \lambda_2 \cdot e^{-\lambda_3 \cdot t}}{(\lambda_1 - \lambda_3) \cdot (\lambda_2 - \lambda_3)} \right]$$

(A17)

We recognize $p_3(t)/N$ where the function $p_3$ has been introduced in (A6a) with $\lambda_1=1/a_1$ , $\lambda_2=1/a_2$ and $\lambda_3=1/a_3$.

#### A.7 Succession of n irreversible chemical reactions

**Case n=1**

We consider a first-order irreversible decomposition chemical reaction, during which the reagent R is transformed into product X as:

$$R \xrightarrow{\upsilon} X$$

where $\upsilon$ is the number of R moles that disappears (or X moles that appears) during the average time $\tau$, i.e. $\upsilon$ is the speed of the chemical reaction; $\upsilon$ is a constant homogeneous to the inverse of a time $(\upsilon \propto 1/\tau)$.

The stationary solution of the macroscopic balance equation of X leads classically [4,5] to:

$$\frac{X(t)}{R_0} = \left(1 - e^{-\upsilon \cdot t}\right) \qquad \text{(A18)}$$

where $R_0$ is the number of moles of reagent present at the beginning of the reaction (t=0).

Equation (A18), which is equivalent to equations (A4a) and (A15), provides the temporal proportion (or average probability) of the reagent transformed into a product during an irreversible chemical reaction.

**Case n=2**

We consider a succession of 2nd order irreversible decomposition chemical reactions, during which the reagent R is transformed into product X through an intermediate state named I**,** where I is both the product of the 1st reaction and the reagent of the second:

$$R \xrightarrow{\upsilon_1} I \xrightarrow{\upsilon_2} X$$

where $\upsilon_1$ and $\upsilon_2$ are the speeds of the 2 successive reactions ; $\upsilon_1$ and $\upsilon_2$ are two constants homogeneous to the inverse of a time, checking $\upsilon_1 \neq \upsilon_2$.

We obtain a system of 3 equations with 3 unknowns, whose resolution leads classically [4] to :

$$\frac{X(t)}{R_0} = \left(1 - \frac{\upsilon_2}{\upsilon_2 - \upsilon_1} \cdot e^{-\upsilon_1 \cdot t} - \frac{\upsilon_1}{\upsilon_1 - \upsilon_2} \cdot e^{-\upsilon_2 \cdot t}\right) \qquad \text{(A19)}$$

where $R_0$ is the number of moles of reagent present at the beginning of the reaction (t=0).

Equation (A19) which is equivalent to equations (A5a) and (A16) provides the temporal proportion (or average probability) of reagent randomly transformed into product during a succession of irreversible chemical reactions.

**Case n=3**

We consider a succession of 3[st] order irreversible decomposition chemical reactions, during which the reagent R is transformed into product X by passing through 2 intermediate states named $I_1$ and $I_2$:

$$R \xrightarrow{\upsilon_1} I_1 \xrightarrow{\upsilon_2} I_2 \xrightarrow{\upsilon_3} X$$

where $\upsilon_1$, $\upsilon_2$ and $\upsilon_3$ are the velocities of the 3 successive reactions, 3 constants homogeneous at the inverse of a time, checking $\upsilon_1 \neq \upsilon_2$, $\upsilon_1 \neq \upsilon_3$ and $\upsilon_2 \neq \upsilon_3$ .

We obtain a system of 4 equations with 4 unknowns whose solution is equivalent to equations (A6a) and (A17). This solution provides the temporal proportion (or average probability) of reagent randomly transformed into product during a succession of irreversible chemical reactions.

**General case**

An obvious link appears between average probabilities relating to successive events occurring at random in time and the kinetic equations of cascading chemical reactions, irreversible and entropy-producing reactions.

By recurrence, we calculate the density of the sum of n independent random variables, each following an exponential law. We deduce from this the probability of n successive random events occurring, which is equivalent to the proportion (or average probability) over time of reagent transformed into product during a succession of $n^{th}$ order irreversible decomposition chemical reactions.

**Notes**

The solution is identical regardless of the order of the n reactions.

The functions $p_1$, $p_2$, $p_3$, $p_4$ and $p_5$ (Fig A1a) whose equations were presented in paragraph A.3 are the solutions of the kinetic equations of one, two, three, four and five successive irreversible reaction(s), respectively.

By reasoning by recurrence following the approximations conducted in (A4b), (A5b) and (A6b) with respect to the functions $p_1(x)$, $p_2(x)$ and $p_3(x)$, respectively, we verify with the expansion to the $n^{th}$ order for the exponential function given in (A3):

$$x \to 0 \quad \Rightarrow \quad p_n(x) \to \frac{x^n}{(n!) \cdot \prod_{i=1}^{n} a_i} \qquad \text{(A20)}$$

where $a_i$ is a constant belonging to $R_+^*$, homogeneous to a time and verifying $a_i \neq a_j$ (i=1,2,…,n ; j=1,2,…, n with i≠j).

The approximation (A20) gives the probability of occurrence for the start (t close to 0) of n successive chemical reactions where each constant $a_i$ is the inverse of the speed of reaction n° i. We observe that at the start of a succession of n reactions, the curve representing the probability of this event and approached by (A21) is flattened when n increases, a phenomenon similar to a time delay (Fig A1b).

#### A.8 Succession of n irreversible chemical reactions where the number of steps is unknown

Many of the mechanisms of the cross-bridge cycle during the implementation of the Working Stroke or during the Recory Stroke remain misunderstood, and as a result, the exact number of reactions involved remains unknown. To do this, we adopt the following method, deduced from the results obtained previously.

The cross-bridge cycle is considered as a set of characteristic states of the myosin head. We pass from one state to another by a global event (G) made up of $n_r$ elementary events occurring successively at random, equivalent to $n_r$ successive irreversible reactions ($n_r$>2).

The entire chain of events leading to the global event G is modeled by the 2 successive random events $K_1$ and $K_2$ representing the 2 irreversible reactions with the 2 slowest reaction rates that are arranged in the last 2 steps, $n_{r-1}$ and $n_r$, since the resulting kinetics is independent of the reaction order. The solution providing the average probability of occurrence of G is identified with that given in (A19), and to take into account the succession of ($n_r$-2) prior irreversible reactions, we introduce a time lag ($\tau_{preK}$) following the approximation (A20) and the associated remark. So:

$$p_G = \left( 1 - \frac{\tau_{K1} \cdot e^{-\frac{t-\tau_{preK}}{\tau_{K1}}}}{(\tau_{K1} - \tau_{K2})} - \frac{\tau_{K2} \cdot e^{-\frac{t-\tau_{preK}}{\tau_{K2}}}}{(\tau_{K2} - \tau_{K1})} \right) \cdot \mathbf{1}_{[\tau_{preK};+\infty[}(t) \qquad \text{(A21)}$$

where $\tau_{K1}$ and $\tau_{K2}$ are 2 time constants expressing the respective mean durations of the 2 slowest of the $n_r$ successive irreversible reactions; $\tau_{preK}$ is the time delay necessary to set up these 2 reactions, a parameter calculated from the sum of the time constants of the ($n_r$-2) other irreversible reactions (if $n_r$ > 2), i.e. $\tau_{preK} = \sum_{i=1}^{n_{r-2}} a_i$ .

The demonstration of (A21) is trivial. In Fig A1b are presented 3 approximations modelled according to (A21) for 3, 4 and 5 successive reactions with time constants identical to those of Fig A1a.

We note:

if $n_r$ = 1, (A18) is equivalent to (A21) with $\tau_{K2}$ = 0 and $\tau_{preK}$ =0

if $n_r$ = 2, (A19) is equivalent to (A21) with $\tau_{preK}$ =0

##### A.9 Conclusion

Any sequence of n irreversible chemical reactions, with n integer greater than or equal to 1, can be modelled by the time equation (A21).
