## Supplementary Chapter. Mathematization of the cross-bridge cycleComputer Programs for calculating and plotting Force/Velocity relationships for "Mechanical model of muscle contraction. 1. Force-velocity relationship"

### S1.B Supplementary Chapter of Paper 1

#### Mathematization of the cross-bridge cycle

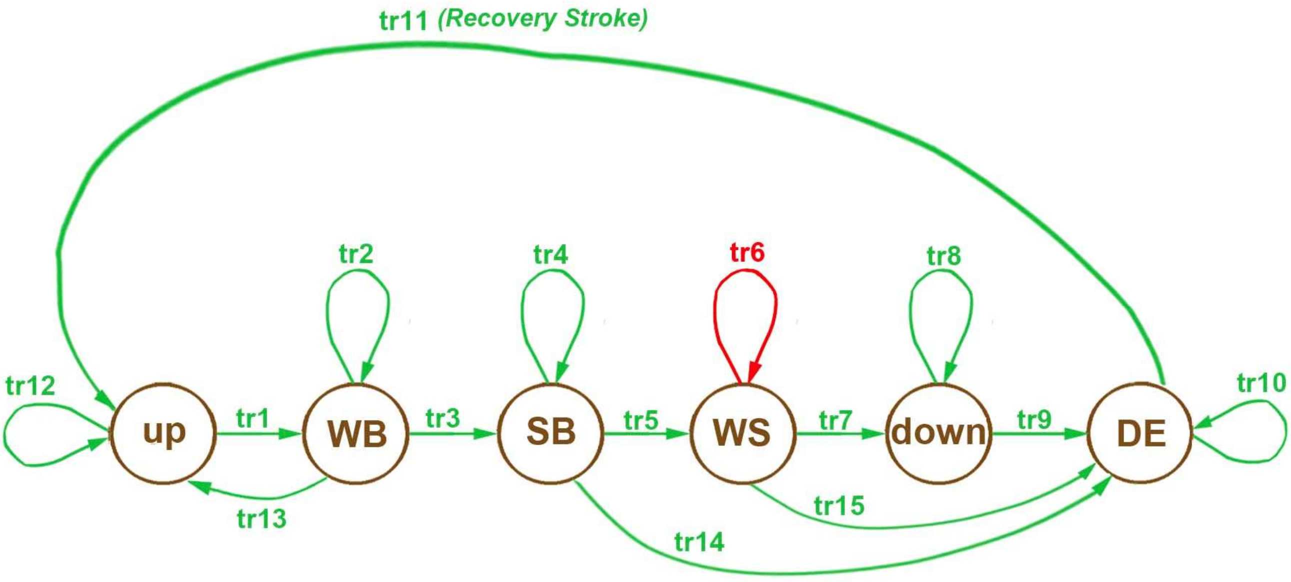


**Fig B1. Graph of the mechanical states and state transitions relative to a myosin head during the cross-bridge cycle.**

**The 6 brown circles represent 6 mechanical states of a myosin head.**

**The 15 green and red arrows are state transitions.**

**The green arrows indicate chemical reactions of entropic origin.**

**The red arrow corresponding to WS Transition n° 6 indicates a mechanically modelable intra-state evolution.**

The series of events occurring to a myosin head analyzed in this appendix is consistent with the conventional description of the cross-bridge cycle [1,2] or the Lymn-Taylor cycle [3], *i.e.* the cycle of chemical-mechanical interactions of the myosin heads with actin filaments, several fundamental mechanisms of which are, to date, not elucidated.

#### B.1 The myosin head as a mechanical object

A myosin II head is modelled by 3 rigid segments [2], articulated between them: the motor domain (S1a), the lever (S1b) and the rod (S2). Six mechanical states related to a myosin head (Fig B1) are traditionally distinguished during the cross-bridge cycle [2,4,5,6,7,8]:

**Up state** (also called as pre-powerstroke, postrecovery, or M**) characterized by the $\theta_{up}$ angle of lever S1b after "rearmament" with Recovery Stroke. The term "rearmament" means "regeneration of the elastic potential energy from which the motor-moment derives".

**WB state** (Weak Binding): S1a is in the vicinity of an actin molecule to which S1a is likely to bind strongly. The word "weak" means that thermal shocks from surrounding molecules can break this bond.

**SB state** (Strong Binding): S1a is recessed in an actin molecule; the term "strong" means that thermal shocks against S1a and S1b cannot easily break the bond.

**WS state** (Working Stroke) is characterized by 4 conditions: (1) the rigidity of S1a, S1b and S2, (2) the state SB, (3) a motor-moment exerted on S1b, a moment which induces a traction of the myosin filament via S2, (4) the displacement of S1b in a fixed plane, the orientation of S1b in this plane being defined with the angle $\theta$ bounded by the two limits $\theta_{down}$ and $\theta_{up}$, parameters defined with equality (8) established in the accompanying Paper 2 and whose values are calculated in (11) and (12).

**Down state** (also called as post-powerstroke, prerecovery, like-rigor or M*) characterized by the angle $\theta_{down}$ of lever S1b. This state closes the WS state with S1a still bound and S1b no longer exerting tensile force on the myosin filament, except for phase 1 of a length step (see paragraphs I.5 and I.6 of Supplement S4.I to accompanying Paper 4).

**DE state** (DEtached): S1a is not weakly or strongly bound to an actin molecule and the angular position of lever S1b is not $\theta_{up}$.

These 6 states are linked by 15 state(s) transitions, 14 of a random nature because produced by chemical reactions of entropic origin (Fig B1; green arrows), and one of a deterministic nature because mechanically modelable (Fig B1; red arrow). The 6 intra-state transitions n° 2, 4, 6, 8, 10 and 12 indicate that a head can remain a variable time in the respective corresponding state WB, SB, WS, down, DE and up, and possibly evolve in this state during the cross-bridge cycle.

#### B.2 The myosin head as a chemical reactor

##### *Hypothesis 1 of irreversibility for reactions localized in the myosin head*

Under physiological conditions, the thermodynamic tables give the following range of values for the variation of free enthalpy of ATP hydrolysis ($\Delta G_{ATP}$):

$$\Delta G_{ATP} \in [-40 \text{ kJ.mole}^{-1} ; -50 \text{ kJ.mole}^{-1}]$$

A chemical reaction is considered irreversible as soon as its variation in free enthalpy is less than -30 kJ.mol$^{-1}$. Following this observation, we formulate hypothesis 1: any transition of state(s) leading to or leaving the WS state consists of a succession of 1$^{st}$ order irreversible chemical reactions.

Hypothesis 1 is illustrated in Fig B1 where all states are linked by single arrows, except for the up and WB states joined by reversible transitions 1 and 13.

The justification for the irreversibility hypothesis is to be read in D. Chowdhury's article [9]. The irreversibility of some stages of the cycle has been described [7,10,11,12,13,14,15,16,17]. However, for many physiologists, the reversibility of the reactions that constitute the cross-bridge cycle remains the rule [3,18,19,20,21,22].

##### *Probability of achieving a mechanical state of the cross-bridge cycle*

Hypothesis 1 mentions that the transitions leading to and leaving the WS state are irreversible and concerns the 5 states SB, WS, down, DE and up. As demonstrated in Supplement S1.A, the probability ($p_G$) of one of these 5 states occurring after a global event named G, composed of a succession of $n_r$ random events, is calculated exactly as the average proportion of a chemical product resulting from a cascade of $n_r$ irreversible reactions and this probability is reducible to the expression (A21) reproduced below:

$$p_G = \left( 1 - \frac{\tau_{K1} \cdot e^{-\frac{t-\tau_{preK}}{\tau_{K1}}}}{(\tau_{K1} - \tau_{K2})} - \frac{\tau_{K2} \cdot e^{-\frac{t-\tau_{preK}}{\tau_{K2}}}}{(\tau_{K2} - \tau_{K1})} \right) \cdot \mathbb{1}_{[\tau_{preK};+\infty[}(t) \qquad \text{(B1a)}$$

where $\tau_{K1}$ and $\tau_{K2}$ are two time constants expressing the mean duration of the two slowest reactions among the $n_r$ successive irreversible reactions; $\tau_{preK}$ is the time necessary to implement these two reactions, a parameter calculated from the sum of the time constants of the ($n_r$-2) other irreversible reactions (if $n_r > 2$); $\mathbb{1}$ is the indicator function defined in (A2b) in the Supplement S1.A.

We distinguish the particular case where a reaction among the $n_r$ successive reactions is significantly slower than the ($n_r$ -1) others. Equation (A21) is simplified in this case and is written:

$$p_K(t) \approx 1 - e^{-\frac{t-\tau_{preK}}{\tau_K}} \cdot \mathbf{1}_{[\tau_{preK};+\infty[}(t) \qquad \text{(B1b)}$$

where $\tau_K$ is the characteristic time constant of the K slowest event leading to the realisation of the state of the cross-bridge cycle, i.e. the slowest reaction among the $n_r$ reactions ($n_r \geq 1$); $\tau_{preK}$ is the characteristic delay time of the pre-setting of K, i.e. the sum of the time constants of the fastest ($n_r$-1) reactions.

If the global event G consists of a single irreversible reaction, the 2 events G and K are confused and the time equation (B1b) remains compatible with the condition $\tau_{preK}=0$.

#### B.3 Initiations of the Working Stroke

Three circumstances lead to the realisation of a WS whose occurrence presupposes the fulfillment of 4 conditions mentioned in paragraph B.1.

**a/ Slow rise of the tension to the isometric tetanus plateau with the event {startS}**

When a fiber initially at rest is tetanized under isometric conditions, the tension slowly increases to the plateau in a few tens of milliseconds [23,24,25,26,27,28,29].

The event consisting of a slow initiation of a WS is called as {startS}; S for *Slow*.

**b/ Very slow component during the isometric tetanus plateau with the event {startVS}**

The plateau is not always perfect and a slight increase is sometimes observable; see for example Fig 11 in [24], and Figs 3 and 6 in [25].

The event consisting in a very slow initiation of a WS is called as {startVS}; VS for *Very Slow*.

**c/ Fast rise of the tension with the event {startF}**

This event concerns the rapid rise observed during phase 2 of a length step with a rise time of about one millisecond [24,26,30,31,32,33,34].

The event consisting in a fast initiation of a WS is called as {startF}; F for *Fast*.

##### *Hypothesis 2*

The initiation event leading to the WS state is called as {WSstart}.

Assumption 2 is enonced: {WSstart} occurs when the θ angle of the lever belongs to the $\delta\theta_{Max}$ interval delimited by $\theta_{down}$ and $\theta_{up}$; {WSstart} is declined according to the 3 modes described above, namely:

| | |
|---|---|
| {startF} | global event related to a fast initiation |
| {startS} | global event related to a slow initiation |
| {startVS} | global event related to a very slow initiation |

Otherwise formulated:

$$\{\text{WSstart}\} \equiv \{\text{startF}\} \cup \{\text{startS}\} \cup \{\text{startVS}\}$$

There are 2 cases depending on whether the {startF} event is feasible or not.

##### *Case 1 where the occurrence of {startF} is possible*

{WSstart} is the combination of the 3 global events {startF}, {startS} and {startVS} and as a myosin head initiating a WS uses only one of the 3 modes, these 3 events are disjoined 2 to 2, that is:

$$P_{WS1}(t) = p_F(t) + p_S(t) + p_{VS}(t) \qquad \text{(B3)}$$

where $P_{WS1}(t)$, $p_F(t)$, $p_S(t)$, $p_{VS}(t)$ are the instantaneous probabilities of occurrence of {WSstart}, {startF}, {startS} and {startVS}, respectively, in Case 1 present.

After an infinite time, i.e. after a few tens of milliseconds corresponding to the arrival of the isometric tetanus plateau, these 4 probabilities tend towards 4 constants defined below:

$$p_F(t) \rightarrow p_{startF} \qquad \text{(B4a)}$$

$$p_S(t) \rightarrow p_{startS} \qquad \text{(B4b)}$$

$$p_{VS}(t) \rightarrow p_{startVS} \qquad \text{(B4c)}$$

$$P_{WS1}(t) \rightarrow (p_{startF} + p_{startS} + p_{startVS}) = 1 \qquad \text{(B4d)}$$

Following hypotheses 1 and 2, we assume that each of the 3 modes is a global event whose occurrence probability is given in (B1b). Taking into account the proportions given in (B4a), (B4b) and (B4c), we pose:

$$p_F(t) = p_{startF} \cdot \left(1 - e^{-\frac{t}{\tau_{startF}}}\right) \qquad \text{(B5a)}$$

$$p_S(t) = p_{startS} \cdot \left(1 - e^{-\frac{t-\tau_{preS}}{\tau_{startS}}}\right) \cdot \mathbf{1}_{[\tau_{preS};+\infty]}(t) \qquad \text{(B5b)}$$

$$p_{VS}(t) = p_{startVS} \cdot \left(1 - e^{-\frac{t-\tau_{preVS}}{\tau_{startVS}}}\right) \cdot \mathbf{1}_{[\tau_{preVS};+\infty]}(t) \qquad \text{(B5c)}$$

The characteristics of each mode are detailed in Table B1.

With (B4d) and the 3 previous time relations, equation (B3) is reformulated:

$$P_{WS1}(t) = p_{startF} \cdot \left(1 - e^{-\frac{t}{\tau_{startF}}}\right) + p_{startS} \cdot \left(1 - e^{-\frac{t-\tau_{preS}}{\tau_{startS}}}\right) + \left(1 - p_{startF} - p_{startS}\right) \cdot \left(1 - e^{-\frac{t-\tau_{preVS}}{\tau_{startVS}}}\right) \qquad \text{(B6)}$$

**Table B1. Characteristics of the global events involved in the tension rise after phase 1 of a length step and in the Force/Velocity relationship.**

| Event {K} | Transition(s) (N° displayed in Fig B1) | $\tau_{preK}$ | Values* (ms) | $\tau_K$ | Values* (ms) |
|---|---|---|---|---|---|
| {SB} | 3 | $\tau_{preSB}$ | 2 à 3 | $\tau_{SB}$ | 6 à 8 |
| {startF} | 5 | $\tau_{preF}$ | 0 | $\tau_{startF}$ | 0.6 à 0.8 |
| {startS} | 1+2+3+4+5 | $\tau_{preS}$ | 2 à 4 | $\tau_{startS}$ | 25 à 30 |
| {startVS} | 11+12+1+2+3+4+5 | $\tau_{preVS}$ | 50 à 80 | $\tau_{startVS}$ | 70 à 130 |
| {FastDE} | 9 | $\tau_{preFDE}$ | ~1 | $\tau_{FDE}$ | 3 à 5 |
| {SlowDE} | 15 | $\tau_{preSDE}$ | 5 à 7 | $\tau_{SDE}$ | 15 à 20 |

(*) Data relating to single fibers isolated from the *tibialis anterior* muscle of two species of frogs, *rana Temporaria* and *rana Esculenta*, under standard conditions with a temperature between 0° and 5°C.

***Case 2 where the occurrence of {startF} is impossible***

In this case {WSstart} is the combination of the 2 global events {startS} and {startVS} according to:

$$P_{WS2}(t) = p_{S2}(t) + p_{VS2}(t) \tag{B3}$$

where $P_{WS2}(t)$, $p_{S2}(t)$ and $p_{VS2}(t)$ are the instantaneous probabilities of realisation of {WSstart}, {startS} and {startVS} in Case 2.

After an infinite time, i.e. after a few tens of ms in reality, these 3 probabilities will tend towards 3 constants defined below:

$$p_{S2}(t) \quad \rightarrow \quad p_{startS2} \tag{B8a}$$

$$p_{VS2}(t) \quad \rightarrow \quad p_{startVS2} \tag{B8b}$$

$$P_{WS2}(t) \quad \rightarrow \quad (p_{startS2} + p_{startVS2})=1 \tag{B8c}$$

Logically $p_{startVS}$ is a constant and remains the same between (B4c) and (B8b):

$$p_{startVS2} = p_{startVS} \tag{B9a}$$

It implies with (B4d), (B8c) and (B9a):

$$p_{startS2} = p_{startF} + p_{startS} \tag{B9b}$$

Each of the 2 modes is a global event composed of several transitions with the occurrence probability provided by expression (B1b). The characteristics of the 2 modes presented in Table B1 do not change from those of Case 1, and equation (B7) is reformulated with (B8c), (B9a) and (B9b):

$$P_{WS2}(t) = (p_{startF} + p_{startS}) \cdot \left(1 - e^{-\frac{t-\tau_{preS}}{\tau_{startS}}}\right) + (1 - p_{startF} - p_{startS}) \cdot \left(1 - e^{-\frac{t-\tau_{preVS}}{\tau_{startVS}}}\right) \tag{B10}$$

#### B.4 Preparation of the event {startF} by the event {SB}

The event {startF} where the WS is initiated quickly is only observed for short length steps, i.e. phase 2 occurs only for a shortening of less than 13 nm per hs; see Fig 9 in [26] and Fig 3B in [31]. On the other hand, the rapid rise observed during phase 2 is all the more important as the duration of tetanization before the length step is important; see Fig 7 in [35]. When a length step of 5 nm follows a length step of 2 nm, the rapid rise in tension is a function of the time between the 2 steps; see Fig 2 in [36]. It is also noted that this rapid rise decreases as the fiber shortening velocity increases; see Figs. 3, 4 and 5 in [37].

All these elements suggest that the {startF} event requires a preparatory step before the length step where heads are strongly bound in SB state.

We complete hypothesis 2 by introducing the probability of the {SB} event occurring before the {startF} event according to the expression (M1b):

$$p_{SB}(t) = p_{startF} \cdot \left( 1 - e^{-\frac{t-\tau_{preSB}}{\tau_{SB}}} \right) \tag{B11}$$

where $p_{startF}$ is the maximum proportion of heads likely to initiate a WS in rapid mode during the isometric tetanus plateau, proportion defined in (B4a); $\tau_{SB}$ is the time constant of the slowest reaction leading to the SB state; $\tau_{preSB}$ is the occurrence delay of this reaction.

**Note:** {SB} and {startF} events are independent since a mechanical action of shortening is required for {startF} to occur after {SB}.

The {SB} event only concerns {startF}. For {startS} and {startVS}, event {SB} is included in these (Table B1). The following inequalities should therefore be checked:

$$\tau_{preSB} < \tau_{preS} < \tau_{preVS}$$

$$\tau_{SB} < \tau_{startS} < \tau_{startVS}$$

#### B.5 Realisation of the DE state according to the WS state

The DE state concerns the detachment of the myosin head from the actin molecule to which it was strongly bound. At the output of the WS state, two transitions to the DE state appear in Fig B1 associated with 2 events:

1/ A fast detachment named {FastDE}: the angle $\theta$ reaches the $\theta_{down}$ limit characterizing the down state with the transition n° 7, then the detachment itself is done in a few milliseconds with the transition n° 9. This event is postulated by several researchers [30,36]. It is studied in the accompanying Papers 4 and 5 with respect to phases 1, 2 and 3 of a length step.

2/ A slow detachment named {SlowDE}: when the angle $\theta$ is between $\theta_T$ and $\theta_{down}$, the moment exerted is low compared to the maximum value of the motor-moment. As a result, the linking actions are reduced which facilitates the exit of the SB state following concomitant shocks of thermal origin (see theoretical calculations in Supplement S2.C of accompanying Paper 2). This event, which takes place as transition n° 15 (Fig B1), has been speculated by several researchers [15,22,38,39].

We assume hypothesis 3: the global event leading to the DE state is broken down into 2 separate events:

{FastDE} event occurring quickly from the down state, i.e. $\theta \leq \theta_{down}$

{SlowDE} slow event from the WS state, i.e. $\theta_{down} \leq \theta \leq \theta_T$

Inequalities are valid in a half-sarcomere on the right and are reversed in a half-sarcomere on the left. Following hypotheses 1 and 3, we assume that each of the 2 modes is a global event whose probability of occurrence is provided by (B1a) or (B1b). The 2 events {FastDE} and {SlowDE} do not concern the same heads and are incompatible.

#### B.6 Detachment in fast mode

The fast detachment follows the shortening by a length step. The occurrence probability of the {FastDE} event ($P_{FDE}$) is calculated according to (B1b):

$$P_{FDE}(t) = \left(1 - e^{-\frac{t-\tau_{preFDE}}{\tau_{FDE}}}\right) . \mathbf{1}_{[-\delta X_{Max};-\delta X_{z1}]}(\Delta X) \quad \text{(B12)}$$

where $\tau_{preFDE}$ is the delay for setting up {FastDE}; $\tau_{FDE}$ is the time constant of {FastDE}; $\delta X_{z1}$ is a linear range defined in paragraph I.6 of Supplement S4.I to accompanying Paper 4.

An order of magnitude of the $\tau_{preFDE}$ and $\tau_{FDE}$ characteristics are given in Table B1.

#### B.7 Detachment in slow mode

This event only concerns WS heads whose lever has an angular position $\theta$ varying between $\theta_T$ and $\theta_{down}$. However, depending on the value of the length step, the WS state of these heads has different origins.

**Option 1 where the WS state is present before the length step**

In this option, the probability of the event occurring ($P_{SDE\_T}$ with "T" for Tetanus) is calculated according to (B1b):

$$P_{SDE\_T}(t) = \left(1 - e^{-\frac{t-\tau_{preSDE}}{\tau_{SDE}}}\right) \quad \text{(B13)}$$

where $\tau_{SDE}$ is the time constant of {SlowDE}; $\tau_{preSDE}$ is the occurence delay necessary for the occurrence of {SlowDE}.

The $\tau_{SDE}$ and $\tau_{preSDE}$ characteristics are given in Table B1.

**Option 2 where the WS state is initiated after the length step**

As previously studied, the occurrence of the global event {WSstart} results from 2 cases depending on whether the event {startF} is feasible or not.

***Option 2 with Case 1 where the occurrence of {startF} is possible after the length step***

We pose:

$$P_{SDE\_1}(t) = p_{startF \to SDE}(t) + p_{startS \to SDE}(t) + p_{startVS \to SDE}(t) \tag{B14}$$

where $P_{SDE\_1}$ is the realisation probability of the {SlowDE} event for Option 2 in Case 1; $p_{startF \to SDE}$, $p_{startS \to SDE}$ and $p_{startVS \to SDE}$ are the occurrence probabilities of {SlowDE} when this event follows {startF}, {startS} or {startVS}, respectively, in the Case 1 present.

As a $\tau_{startF}$, the time constant of {startF}, is much lower than $\tau_{SDE}$, the mean duration of {SlowDE}, the probability $p_{startF \to SDE}$ is calculated according to (B1b):

$$p_{startF \to SDE}(t) = p_{startF} \cdot \left( 1 - e^{-\frac{t - \tau_{preSDE}}{\tau_{SDE}}} \right) \tag{B15a}$$

Since $\tau_{startS}$ and $\tau_{startVS}$, the time constants of {startS} and {startVS}, are of the same order of magnitude as $\tau_{SDE}$, the mean duration of {SlowDE}, the formula (B1a) is applied with (B5b) and (B5c) for the respective calculations of $p_{startS \to SDE}$ and $p_{startVS \to SDE}$:

$$p_{startS \to SDE}(t) = p_{startS} \cdot \left( 1 - \frac{\tau_{startS} \cdot e^{-\frac{t-(\tau_{preS}+\tau_{preSDE})}{\tau_{startS}}}}{(\tau_{startS} - \tau_{SDE})} - \frac{\tau_{SDE} \cdot e^{-\frac{t-(\tau_{preS}+\tau_{preSDE})}{\tau_{SDE}}}}{(\tau_{SDE} - \tau_{startS})} \right) \tag{B15b}$$

$$p_{startVS \to SDE}(t) = (1 - p_{startF} - p_{startS}) \cdot \left( 1 - \frac{\tau_{startVS} \cdot e^{-\frac{t-(\tau_{preVS}+\tau_{preSDE})}{\tau_{startVS}}}}{(\tau_{startVS} - \tau_{SDE})} - \frac{\tau_{SDE} \cdot e^{-\frac{t-(\tau_{preVS}+\tau_{preSDE})}{\tau_{SDE}}}}{(\tau_{SDE} - \tau_{startVS})} \right) \tag{B15c}$$

The temporal parameters for all these events are given in Table B1 with their respective values for fast frog fibers.

***Option 2 with Case 2 where the occurrence of {startF} is impossible after the length step***

We pose:

$$P_{SDE\_2}(t) = p_{startS2\rightarrow SDE}(t) + p_{startVS2\rightarrow SDE}(t) \qquad \text{(B16)}$$

where $P_{SDE\_2}(t)$ is the realisation probability of the {SlowDE} event for Option 2 in Case 2; $p_{startS2\rightarrow SDE}(t)$ and $p_{startVS2\rightarrow SDE}(t)$ are the occurrence probabilities of the {SlowDE} event when this event succeeds either {startS} or {startVS}, respectively, in the Case 2 here.

With (B9a) and (B9b), these probabilities are formulated according to (B1a):

$$p_{startS2\rightarrow SDE}(t) = \left(p_{startF} + p_{startS}\right)\cdot\left(1 - \frac{\tau_{startS}\cdot e^{-\frac{t-\tau_{preS}-\tau_{preSDE}}{\tau_{startS}}}}{\left(\tau_{startS}-\tau_{SDE}\right)} - \frac{\tau_{SDE}\cdot e^{-\frac{t-\tau_{preS}-\tau_{preSDE}}{\tau_{SDE}}}}{\left(\tau_{SDE}-\tau_{startS}\right)}\right) \qquad \text{(B17a)}$$

$$p_{startVS2\rightarrow SDE}(t) = p_{startVS\rightarrow SDE}(t) \qquad \text{(B17b)}$$

The characteristics of each mode are detailed in Table B1.
