## Supplementary material for "Mechanical model of muscle contraction. 1. Force-velocity relationship": Computer Programs for calculating and plotting Force/Velocity relationships

### CP1 Computer programs for calculating and plotting Force/Velocity relationships

The curves on the computer screen are done by calling the "Sub AA_FVtouch()" routine.

Once the data have been recovered (See values displayed in the Tables of Paper 1 and in the Excel sheets of Supplement DA1), the experimental points from published articles are plotted and the theoretical tension values from model equations are calculated for the corresponding velocities according by running the subroutine "AA_FVtouch_calculs_OK_dec_2018"

A regression is performed between experimental and theoretical tensions.

Then we draw on 200 values the F/V curves according to the equations of the model by using again the subroutine "AA_FVtouch_calculs_OK_dec_2018" (Fig CP1.1).

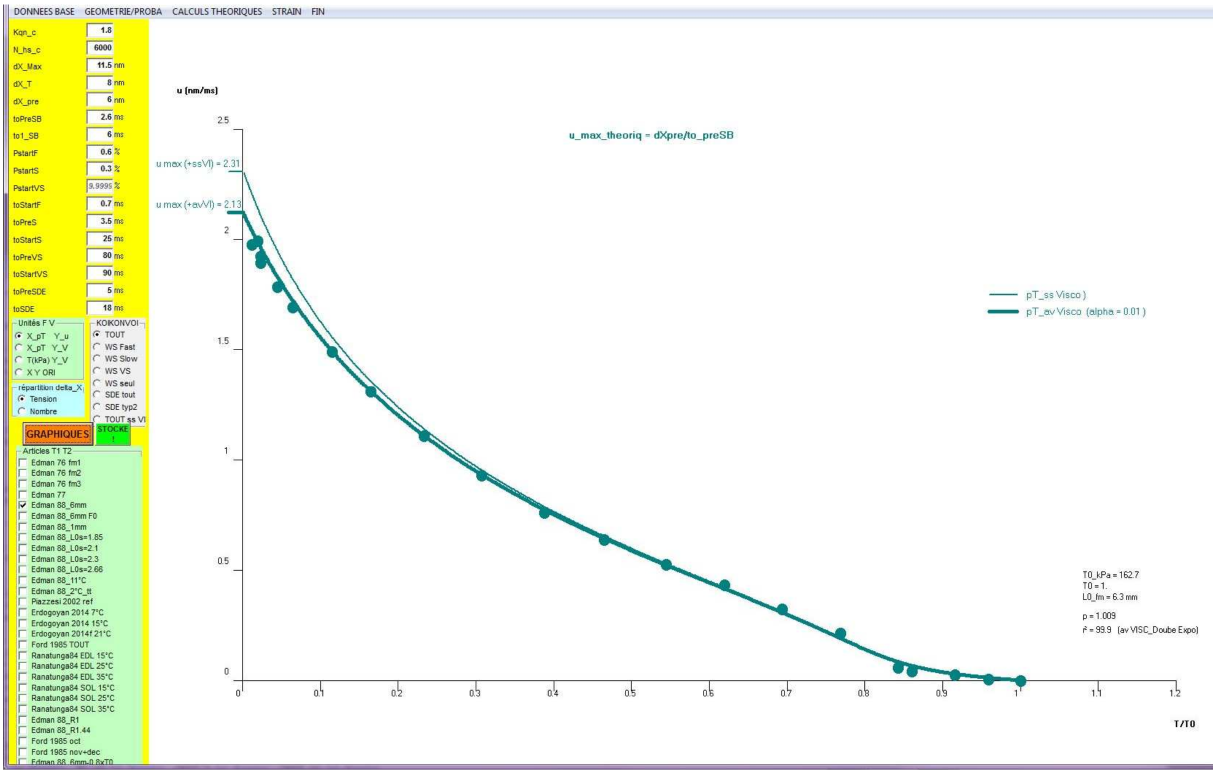


**Fig CP1.1 Screenshot after starting the "Sub AA_FVtouch()" routine**

### Sub AA_FVtouch()

```
Dim T0_etoile As Single, T0_kPa As Single
'Rappel les caluls pour avoir toutes les variables (u V TpT) fait dans  FORM_LOAD en applelant
```

**SUB_AA_FVtouch_stocke_fm 25**

```
txV = "ss_T0_WS"
'txV = "av_T0_WS"

'exposant_visc = 0 ' regression sans viscosité
exposant_visc = 1 ' regression avec viscosité

O.Cls
'on sait combien il y a de fm traitées
If N_fm = 1 Then 'c'est bon on travaille
    KqnZ1 = Val(TAM(0).Text) 'ne sert à rien
    N_hs = Val(TAM(1).Text) 'sert

    L_WS = Val(TAM(2).Text) 'dX_Max = 12 nm
    dX_T = Val(TAM(3).Text) 'dX_T =8.5 nm
    aX = dX_T 'pour etre tranquille
    L_preWS = Val(TAM(4).Text) 'dX_pre = 5.5 nm

    t_preSB = Val(TAM(5).Text) '0 ms
    to1_SB = Val(TAM(6).Text) '6 ms

    P_startF = Val(TAM(7).Text)
    P_startS = Val(TAM(8).Text)
    'petite verif
    P_startH = P_startF + P_startS
    If P_startH > 1 Then
        vd = MsgBox("OBLIGATOIR : p_startF + p_startS <= 1", 0, "GAFFE !")
        Exit Sub
    End If
    P_startVS = 1 - P_startH
    TAM(9).Text = Format(P_startVS)

    to_startF = Val(TAM(10).Text)
    t_preS = Val(TAM(11).Text)
    to1_startS = Val(TAM(12).Text)
    t_preVS = Val(TAM(13).Text)  '70-150 ms
    to1_startVS = Val(TAM(14).Text) '100 ms

    t_pre_SlowDE = Val(TAM(15).Text)  't>10 ms ?
    to_SlowDE = Val(TAM(16).Text)     '= to_startS = 30 ms ?

    'calcul_qoK (KqnZ1) '1 / 8.01) * Log(2800000 / Kqn) '1.9
    qZ1 = Val(TAM(17).Text) ' = Format(qOK, "#.###")

    FVto.Seek "=", zfm, -1: If FVto.NoMatch Then Stop
    model_ViSC = FVto("model")
```

**AA_FVtouch_recup_data_VISCO**

```
End If


Np4 = 100
e_pt = IIf(FraX.Tag = 0, 15, 25) '15 '12 '*
e_tt = IIf(FraX.Tag = 0, 4, 7) '5 '* IIf(FraX.Tag = 0, 1, 2)
```

```
'pb d'echelle
u_max = 0
For t = 0 To N_articles - 1
   If ChT1T2(t) = 1 Then 'c'est ok
       'en 1 pour temp etudie ensemble a partir du T0 de 0°C
        zfm = ChT1T2(t).Tag - 900
        'N_fm = N_fm + 1:
        FVto.Seek "=", zfm, 1: If FVto.NoMatch Then Stop
        If u_max < FVto("u") Then u_max = FVto("u")
    End If
Next
v = Int(u_max / 2.5) + 1 ' par defaut
'v = 1.6 'pour les 4 traces de Viscosité si 15%
'v = 3 'pour les 4 traces de Viscosité si 20%
'v = Int(u_max / 2.5) + 2 '2 pour EDL ds Ranantunga 84
'v = Int(u_max / 2.5) + 4 '4 pour SOL ds Ranantunga 84

'X0=-5: larg=110
If FraX.Tag = 0 Then
   If (zfm >= 8 And zfm <= 13) Or (zfm >= 19 And zfm <= 24) Then
      impscale -10, Y0 + 140, X0 + 200, Y0 '+ 5 '14
   Else
    impscale -10, Y0 + 140, X0 + 115, Y0 '+ 5 '14
   End If

Else 'PN en fonction de pT
   impscale -10, Y0 + 140, X0 + 215, Y0 '+ 5 '14
End If

If FraX.Tag = 0 Then 'Tension
    Select Case Frtyp.Tag
     Case 1 'X=pT Y=u
          zY1 = "pT": zY2 = "u"
     Case 2 'X=pT Y=u
          zY1 = "pT": zY2 = "V"
     Case 3 'X=T Y=u
          zY1 = "T_kPa": zY2 = "V"
     Case 4 'X Y ori
         'if n_fm>1 then pbroblme?
         FVto.Seek "=", zfm, -1: If FVto.NoMatch Then Stop
         zY1 = FVto("ori_T"): zY2 = FVto("ori_V")
    End Select
Else 'NOmbre
    zY1 = "pT": zY2 = "pN"
End If

  Select Case zY1 'TENSION
     Case "pT": vx = "T/T0": xn = 0: xx = IIf(zY2 = "pN", 1, 1.2): xt = 0.1
     Case "T_Nmm2": vx = "T (N/mm²)": xn = 0: xx = 0.4: xt = 0.05
     Case "T_kPa": vx = "T (kPa)": xn = 0: xx = 250: xt = 25
     Case "T_mN": vx = "T (mN)": xn = 0: xx = 6: xt = 0.5
     Case Else: Stop
    End Select
    zY4 = IIf(zY1 = "pT", "p", "")

  Select Case zY2 'VITESSE ou N
     Case "u": vy = "u (nm/ms)": yn = 0:  yt = 0.5: yx = 2.5 * v '2.5
     Case "V": vy = "V (mm/s)": yn = 0:  yt = 5: yx = 25 * v '35 pour edman 76 77
```

```
    Case "V_L_s": vy = "V (L/s)": yn = 0:  yt = 0.5: yx = 2.5 * v
    Case "pN": vy = "N/N0": yn = 0: yx = 1.1: yt = 0.1
    Case Else: Stop
   End Select

impenvi
If FraX.Tag = 1 Then
   O.DrawStyle = 2
   yav = (1 - yn) * 100 / (yx - yn)
   O.Line (0, yav)-(100, yav), QBColor(3)
   O.DrawStyle = 0
End If
'********* etap 1 : trace ds points enregistrés ds table_T1T2

  O.DrawMode = 13

   For t = 0 To N_articles - 1
    If ChT1T2(t) = 1 Then 'c'est ok
      'en 1 pour temp etudie ensemble a partir du T0 de 0°C
      zfm = ChT1T2(t).Tag - 900
      'N_fm = N_fm + 1:
      FVto.Seek "=", zfm, -1: If FVto.NoMatch Then Stop
      T0m1 = IIf(FraX.Tag = 0, FVto(zY1), 1)
      L0 = FVto("L0_fm")
      T0_kPa = FVto("T_Kpa")
      If IsNull(FVto("N_hs")) Then 'on ne le fait que la 1ere fois
            vd = MsgBox("ZUT !é", 0, "Faire seule la fm n°" + Format(zfm))
            Exit Sub
      End If
      model_ViSC = FVto("model")
      If N_fm > 1 Then 'N_fm calcul ds chT1T2_click
          N_hs = FVto("N_hs")
           L_WS = FVto("dX_Max")
           dX_T = FVto("dX_T"): aX = dX_T
           L_preWS = FVto("dX_pre")
           t_preSB = FVto("t_preSB")
           to1_SB = FVto("to_SB")
           'to_Relax = FVto("to_Relax")
           P_startF = FVto("P_startF")
           P_startS = FVto("P_startS")
           to_startF = FVto("to_startF")
           P_startVS = 1 - P_startF - P_startS
          't_preS = FVto("t_preS")
          to1_startS = FVto("to_startS")
          t_preVS = FVto("t_preVS")
          to1_startVS = FVto("to_startVS")
          't_pre_FastDE = FVto("t_pre_FastDE")
          'to_FastDE = FVto("to_FastDE")
          t_pre_SlowDE = FVto("t_preSDE")
          to_SlowDE = FVto("to_SDE")
```

**AA_FVtouch_recup_data_VISCO**

```
      End If

   '******************PONITS MESURES et on stocke valeurs theoriques correspondants à u
      sN = 0: sX = 0: sXX = 0: sY = 0: sYY = 0: sXY = 0
      FVto.Seek "=", zfm, 1: If FVto.NoMatch Then Stop
      coul = QBColor(FVto("coul"))
```

```
u_max = FVto("u"): If u_max < 0.1 Then Stop
O.DrawWidth = e_pt
Do While FVto("fm") = zfm

   u = -FVto("u")
   If u < 0 Then 'calul pour les points thoeriques correspondants
     AA_FVtouch_calculs_OK_dec_2018
   Else ' u=0 :isométrie
     xmi = L_WS ^ 2 / (2 * dX_T * Abs(X3))
     pT_F = P_startF * xmi
     pT_S = P_startS * xmi
     pT_VS = (1 - P_startF - P_startS) * xmi
     pT_SDE1 = Xmin ^ 2 / (2 * dX_T * Abs(X3)): pT_SDE_simp1 = pT_SDE1:  pT_SDE_tt = pT_SDE1
     pT_SDE2 = 0
     pT_vi = 0

     pN_F = P_startF * L_WS / dX_T
     pN_S = P_startS * L_WS / dX_T
     pN_VS = (1 - P_startF - P_startS) * L_WS / dX_T
     pN_SDE1 = Abs(Xmin) / dX_T: pN_SDE_simp1 = pN_SDE1:  pN_SDE_tt = pN_SDE1
     pN_SDE2 = 0
   End If

   FVto.Edit
      FVto("pT_tot") = pT_F + pT_S + pT_VS - pT_SDE1 - pT_SDE2
      FVto("diff") = FVto("pT_tot") - FVto("pT") '= (pT_theoriq - pT_mesurée)
      FVto("pT_F") = pT_F
      FVto("pT_S") = pT_S
      FVto("pT_VS") = pT_VS
      FVto("pT_SDE_simpl") = pT_SDE_simp1
      FVto("pT_SDE1") = pT_SDE1
      FVto("pT_SDE2") = pT_SDE2
      FVto("pT_SDE_tt") = pT_SDE_tt
      FVto("pT_vi") = pT_vi
      FVto("pT_tot_avVI") = FVto("pT_tot") - pT_vi
      FVto("diff2") = FVto("pT_tot_avVI") - FVto("pT")

      FVto("pN_tot") = pN_F + pN_S + pN_VS - pN_SDE1 - pN_SDE2
      FVto("pN_F") = pN_F
      FVto("pN_S") = pN_S
      FVto("pN_VS") = pN_VS
      FVto("pN_SDE_simpl") = pN_SDE_simp1
      FVto("pN_SDE1") = pN_SDE1
      FVto("pN_SDE2") = pN_SDE2
      FVto("pN_SDE_tt") = pN_SDE_tt
  FVto.Update


  'regression   avec ou ss vi
  If FraX.Tag = 0 Then
   sN = sN + 1
   sX = sX + FVto("pT"): sXX = sXX + FVto("pT") ^ 2

   SDg = IIf(exposant_visc = 0, FVto("pT_tot"), FVto("pT_tot_avvi"))
   sY = sY + SDg: sYY = sYY + SDg ^ 2
   sXY = sXY + FVto("pT") * SDg
  End If
```

```
          xav = FVto(zY1) 'T
          If FraX.Tag = 0 Then
             yav = FVto(zY2)
          Else
             yav = FVto("pN_tot")
          End If
          xav = (xav - xn) * 100 / (xx - xn)
          yav = (yav - yn) * 100 / (yx - yn)

          O.PSet (xav, yav), coul
          FVto.MoveNext: If FVto.EOF Then Exit Do
       Loop

  '
If FraX.Tag = 0 Then 'on calcule  'r
  'r_regL = (sXY - sX * sY / sN) / Sqr((sXX - sX * sX / sN) * (sYY - sY * sY / sN))'r classqie
  'r_regL = sXY / Sqr(sXX * sYY)  'r pente à zro

 'pente
 ' p_regL = (sXY - sX * sY / sN) / (sXX - sX * sX / sN)
   p_regL = sXY / sXX
   r_regL = 1 - (sYY - 2 * p_regL * sXY + sXX * p_regL ^ 2) / (sYY - sY * sY / sN)

   If Abs(r_regL) > 1 Then Stop 'bizarre
   O.CurrentX = 90: O.CurrentY = 12.5: O.Print "p = " + Format(p_regL, "0.###")
   O.CurrentX = 90: O.CurrentY = 10: O.Print "r² = " + Format(r_regL * 100, "###.##") + "   (" + IIf(exposant_visc = 0, "ss",
"av") + " VISC_" + Choose(model_ViSC + 1, "Doube Expo", "2Droites", "Sigmoide") + ")"

     O.CurrentX = 90: O.CurrentY = 20: O.Print "T0_kPa = " + Format(T0_kPa, "###.#")
     O.CurrentX = 90: O.CurrentY = 18: O.Print "T0 = " + Format(T0m1, "##0.###")
     O.CurrentX = 90: O.CurrentY = 16: O.Print "L0_fm = " + Format(L0, "#0.#") + " mm"

  'ordonné à l'origine
  'Y0_regL = sY / sN - p_regL * sX / sN

End If
   '********** COURBE THEORIQUE continue

      O.FontSize = 10
      Np4 = 200

     '                        1      2       3       4      5      6        7       8
     TRUC = Fresc.Tag '0_TOUT_ss_VI 1_WSfast 2_WSslow 3_WSvery_slow 4_WSseul 5_SDEtout 6_SDE_typ2
7_toutssVI  TOUT ss VI
     For k = IIf(TRUC = 0, 7, TRUC) To IIf(TRUC = 0, 8, TRUC)
     'For k = IIf(TRUC = 0, 7, TRUC) To IIf(TRUC = 0, 7, TRUC) 'pour les 4/5 traces de la viscosité

     'le point (T0*;u=0) mais en fait corrrspnd à celui decrit dans la litterature on laisse tomber
        If txV = "av_T0_WS" And (k <= 4 Or k >= 7) Then
           O.DrawWidth = 2
          'coul = QBColor(0) '&HFF00&

         'T0
          xap = (T0m1 - xn) * 100 / (xx - xn)
          O.Line (xap, -1)-(xap, -10), coul
          O.CurrentX = xap - 5: O.CurrentY = -5: O.Print zY4 + "T0 =" + Format(T0m1, "##0.##")

         'T0*
```

```
   T0_etoile = L_WS ^ 2 / (2 * dX_T * Abs(X3))
   If zY1 <> "pT" Then T0_etoile = T0_etoile * T0m1
   xap = (T0_etoile - xn) * 100 / (xx - xn)
   O.Line (xap, -1)-(xap, -10), coul
   O.CurrentX = xap + 0.5: O.CurrentY = -3: O.Print zY4 + "T0_WS =" + Format(T0_etoile, "##0.###")
   O.CurrentX = xap + 0.5: O.CurrentY = -5: O.Print zY4 + "T0 =" + Format(T0m1 / T0_etoile, "0.###") + " x " + zY4 +
"T0_WS*"

  End If


  If txV = "av_T0*" And FraX.Tag = 0 And k >= 7 Then 'on trace fin en pointille sans SDE jusqu'à pT0*
    O.DrawWidth = 1
    O.DrawStyle = 2
   xmi = T0_etoile
   O.CurrentX = (xmi - xn) * 100 / (xx - xn)
   O.CurrentY = -yn * 100 / (yx - yn)
   For i = 1 To Np4
      u = -i * u_max / Np4
```

**AA_FVtouch_calculs_OK_dec_2018**

```
      xmi = pT_F + pT_S + pT_VS
      Select Case zY2 'VITESSE
       Case "u": ymi = -u
       Case "V": ymi = -u * N_hs / 1000
       Case "V_L_s": ymi = -u * N_hs / (L0 * 1000)
      End Select

      xap = (xmi * T0m1 - xn) * 100 / (xx - xn)
      If xap < 0 Then Exit For
      yap = (ymi - yn) * 100 / (yx - yn)
    O.Line -(xap, yap), coul
   Next
   O.DrawStyle = 0
  End If

 '---
 '       O.DrawWidth = e_tt '* 5
  'O.DrawWidth = IIf(k = 8, 1, e_tt)
  O.DrawWidth = IIf(k = 7, 1, e_tt)
  'point isométrie (pT0=1 u=0
  If FraX.Tag = 0 Then

   xmi = IIf(k = 4, T0_etoile, T0m1)
   O.CurrentX = (xmi - xn) * 100 / (xx - xn)
   O.CurrentY = -yn * 100 / (yx - yn)
  Else

   O.CurrentX = 100 '(T0m1 - xn) * 100 / (xx - xn)
   O.CurrentY = (1 - yn) * 100 / (yx - yn)
  End If

  Select Case zfm
   'Case 5: u_max = 8: coul = QBColor(10)
   Case 1, 2, 4, 5, 11, 13, 22, 27, 28: u_max = 2.5
   Case 3, 14, 25, 26: u_max = 4
   Case 15, 19, 23: u_max = 10
   Case 16, 17, 20, 21, 24: u_max = 15
  End Select
```

```
 'pour 5trac"s de viscosit
 If exposant_visc = 1 And zfm = 5 Then
   'coul = QBColor(3) 'bleu-vert
   'coul = QBColor(8) 'gris
   'coul = QBColor(10) 'vert
   'coul = QBColor(13) 'mauve
   coul = QBColor(12) 'rouge
   u_max = 8
 End If

 For i = 1 To Np4
   u = -i * u_max / Np4
   AA_FVtouch_calculs_OK_dec_2018

   If FraX.Tag = 0 Then
      '                       1       2          3         4        5        6         7
      'TRUC = Fresc.Tag '0_TOUT 1_WSfast 2_WSslow 3_WSvery_slow 4_WSseul 5_SDEtout 6_SDE_typ2
7_tout+VI

     '    k =          1     2     3 |          4          |    5    |      6      |                  7        |               8
     xmi = Choose(k, pT_F, pT_S, pT_VS, pT_F + pT_S + pT_VS, pT_SDE_tt, pT_SDE1, pT_F + pT_S + pT_VS -
pT_SDE_tt, pT_F + pT_S + pT_VS - pT_SDE_tt - pT_vi)
     'If xmi > 1.01 Then Stop
     Select Case zY2 'VITESSE
      Case "u": ymi = -u
      Case "V": ymi = -u * N_hs / 1000
      Case "V_L_s": ymi = -u * N_hs / (L0 * 1000)
     End Select

    Else 'NOMBRE
     xmi = pT_F + pT_S + pT_VS - pT_SDE1 - IIf(k = 8, pT_vi, 0)

     '    k =          1     2     3 |          4        |    5  |       6      |              7            |               8
     ymi = Choose(k, pN_F, pN_S, pN_VS, pN_F + pN_S + pN_VS, pN_SDE_tt, pN_SDE1, pN_F + pN_S + pN_VS -
pN_SDE_tt, pN_F + pN_S + pN_VS - pN_SDE_tt)
   End If

   xap = (xmi * T0m1 - xn) * 100 / (xx - xn)
   If xap < 0 Then Exit For
   yap = (ymi - yn) * 100 / (yx - yn)
   O.Line -(xap, yap), coul
 Next

If model_ViSC = 10 Then
  O.DrawWidth = 1:  O.DrawStyle = 2
  'u1
  ymi = (u0 - yn) * 100 / (yx - yn)
  O.CurrentX = -1.5: O.CurrentY = ymi + 1.5: O.Print "u1"
  O.Line (0, ymi)-(70, ymi)
  'u2
  ymi = (u2 - yn) * 100 / (yx - yn)
   O.CurrentX = -1.5: O.CurrentY = ymi + 1.5: O.Print "u2"
  O.Line (0, ymi)-(45, ymi)
End If
 'puiisance FV
 'O.DrawWidth = e_tt * 5
```

```
        'FVto.Seek "=", z, 1: If FVto.NoMatch Then Stop

        'Do While FVto("fm") = z
        '   xav = -FVto("V3"): xav = (xav - xn) * 100 / (xx - xn)
            '*****VERSION 1
        '   yav = FVto("pT_tot") * FVto("V3"): yav = (yav - yn) * 100 / (yx - yn)
        '    O.PSet (xav, yav), QBColor(12)


        '   FVto.MoveNext: If FVto.EOF Then Exit Do
        'Loop
    '
    Next 'k

    End If
  Next 'article


End Sub
```

#### Sub AA_FVtouch_recup_data_VISCO()

```
 Select Case model_ViSC
    Case 0 'Double exponentielle
        pT_Vi_sta = 0
        If IsNull(FVto("DE_pT_Vi_sta")) = 0 Then
          pT_Vi_sta = FVto("DE_pT_Vi_sta")
          to1_u = FVto("DE_to1_u")
          to2_u = FVto("DE_to2_u")
          u0 = FVto("DE_u0")
        End If
     Case 1 'Deux Droites
       If IsNull(FVto("D2_pente1")) = 0 Then
          pente1 = FVto("D2_pente1")
          u0 = FVto("D2_u1")
          pente2 = FVto("D2_pente2")
          u2 = FVto("D2_u2")
        End If
     Case 2 'Sigmoide
        pT_Vi_sta = 0
        If IsNull(FVto("S_pT_Vi_sta")) = 0 Then
          pT_Vi_sta = FVto("S_pT_Vi_sta")
          to1_u = FVto("S_to")
          uI = FVto("S_uI")
          u0 = 0.1
        End If
     End Select

End Sub
```

#### Sub AA_FVtouch_calculs_OK_dec_2018()

```
'la vitesse u doit être négative
If u >= 0 Then Stop

X1 = dX_T / 2      'X_up
 X2 = -X1         'X_T
 X3 = X1 - L_WS   'X_down
 Xmin = dX_T - L_WS '-dX_E <0

'******** WS **************

'Evt startF
Xa = P_startF * (1 - Exp((L_preWS + u * t_preSB) / (u * to1_SB))) 'qF(u)
RS1a = (1 - Exp(L_WS / (u * to_startF)))
pN_F = (Xa / dX_T) * (L_WS + u * to_startF * RS1a)
pT_F = (Xa * L_WS / (Abs(X3) * dX_T)) * (L_WS / 2 + u * to_startF + (u * to_startF) ^ 2 * RS1a / L_WS)
'If pT_F < 0 Then pT_F = 0 'Stop
' If FVto("vu") = "F0" Then Stop

'Evt startS
Xb = P_startS + P_startF - Xa 'qS(u)
RS1b = (1 - Exp(L_WS / (u * to1_startS)))
pN_S = (Xb / dX_T) * (L_WS + u * to1_startS * RS1b)
pT_S = (Xb * L_WS / (Abs(X3) * dX_T)) * (L_WS / 2 + u * to1_startS + (u * to1_startS) ^ 2 * RS1b /
L_WS)

'Evt startVS
If Xa + Xb > 1 Then Stop
If Abs(u) * t_preVS < L_WS Then
   X_VS = X1 + u * t_preVS
   If X_VS < X3 Then Stop
   Xc = 1 - P_startF - P_startS
   RS2 = (1 - Exp((X_VS + Abs(X3)) / (u * to1_startVS)))
   pN_VS = (Xc / dX_T) * (X_VS + Abs(X3) + u * to1_startVS * RS2)
   pT_VS = (Xc * (X_VS + Abs(X3)) / (Abs(X3) * dX_T)) * ((X_VS + Abs(X3)) / 2 + u * to1_startVS + (u *
to1_startVS) ^ 2 * RS2 / (X_VS + Abs(X3)))

 Else
   Xc = 0: pN_VS = 0: pT_VS = 0
End If

'******** Slow DE **************
'pN_SDE nbre relatif de tetes se detachant lentement
'pT_SDE tension relative crrée par les tetes se detachant lentementnbre de SDE
If Abs(u) * t_pre_SlowDE < (L_WS - aX) Then
    X_SDE = -aX / 2 + u * t_pre_SlowDE 'X_T= -ax/2
   If X_SDE < -L_WS Then Stop

'type 1
  'qSDE(u)
    MS1a = 1 - Exp(dX_T / (u * to_startF))
    MS1b = 1 - Exp(dX_T / (u * to1_startS))
    MS2 = 0
```

```
   If Xc > 0 Then MS2 = 1 - Exp((dX_T + u * t_preVS) / (u * to1_startVS))

  'pT_SDE1 (mon truc du depart)
   Xd = Xa * MS1a + Xb * MS1b + Xc * MS2
   Rm = (1 - Exp((X_SDE + Abs(X3)) / (u * to_SlowDE)))
   pN_SDE1 = (Xd / dX_T) * (X_SDE + Abs(X3) + u * to_SlowDE * Rm)
   pT_SDE1 = (Xd * (X_SDE + Abs(X3)) / (Abs(X3) * dX_T)) * ((X_SDE + Abs(X3)) / 2 + u * to_SlowDE
+ (u * to_SlowDE) ^ 2 * Rm / (X_SDE + Abs(X3)))

  'pT_SDE_simp (pour simplifier mes formules)
   Xd = Xa + Xb + Xc * MS2
   'Rm = (1 - Exp((X_SDE + Abs(X3)) / (u * to_SlowDE)))
   pN_SDE_simp1 = (Xd / dX_T) * (X_SDE + Abs(X3) + u * to_SlowDE * Rm)
   pT_SDE_simp1 = (Xd * (X_SDE + Abs(X3)) / (Abs(X3) * dX_T)) * ((X_SDE + Abs(X3)) / 2 + u *
to_SlowDE + (u * to_SlowDE) ^ 2 * Rm / (X_SDE + Abs(X3)))

'type 2 Pdp As Single, Qdp As Single, Ddp As Single
 'F-->SDE2
   MS1a = Exp(dX_T / (u * to_startF))
   Xd = Xa * MS1a
   Rm = (1 - Exp((X_SDE + Abs(X3)) / (u * to_SlowDE)))
   pN_SDE2 = (Xd / dX_T) * (X_SDE + Abs(X3) + u * to_SlowDE * Rm)
   pT_SDE2 = (Xd * (X_SDE + Abs(X3)) / (Abs(X3) * dX_T)) * ((X_SDE + Abs(X3)) / 2 + u * to_SlowDE
+ (u * to_SlowDE) ^ 2 * Rm / (X_SDE + Abs(X3)))

  'S-->SDE2
   MS1b = Exp(dX_T / (u * to1_startS))
   Xd = Xb * MS1b
   'Rm = (1 - Exp((X_SDE + Abs(X3)) / (u * to_SlowDE)))
   Ddp = (1 - Exp((X_SDE + Abs(X3)) / (u * to1_startS)))

   Pdp = u * to_SlowDE ^ 2 * Rm / (to_SlowDE - to1_startS)
   Qdp = u * to1_startS ^ 2 * Ddp / (to1_startS - to_SlowDE)

   pN_SDE2 = pN_SDE2 + (Xd / dX_T) * (X_SDE + Abs(X3) + Pdp + Qdp)
   pT_SDE2 = pT_SDE2 + (Xd * (X_SDE + Abs(X3)) / (Abs(X3) * dX_T)) * ((X_SDE + Abs(X3)) / 2 + u *
(to_SlowDE + to1_startS) + (u * to_SlowDE * Pdp + u * to1_startS * Qdp) / (X_SDE + Abs(X3)))

   'VS-->SDE2
   If Xc > 0 Then
     MS2 = Exp((dX_T + u * t_preVS) / (u * to1_startVS))
     Xd = Xc * MS2
     'Rm = (1 - Exp((X_SDE + Abs(X3)) / (u * to_SlowDE)))
     Ddp = (1 - Exp((X_SDE + Abs(X3)) / (u * to1_startVS)))

     Pdp = u * to_SlowDE ^ 2 * Rm / (to_SlowDE - to1_startVS)
     Qdp = u * to1_startVS ^ 2 * Ddp / (to1_startVS - to_SlowDE)

    pN_SDE2 = pN_SDE2 + (Xd / dX_T) * (X_SDE + Abs(X3) + Pdp + Qdp)
    pT_SDE2 = pT_SDE2 + (Xd * (X_SDE + Abs(X3)) / (Abs(X3) * dX_T)) * ((X_SDE + Abs(X3)) / 2 + u *
(to_SlowDE + to1_startVS) + (u * to_SlowDE * Pdp + u * to1_startVS * Qdp) / (X_SDE + Abs(X3)))

  End If
```

```
 'pN_SDE_tt (les onnes formules) à comparer avec pT_SDE1 et pT_SDE_simpl
  pN_SDE_tt = pN_SDE1 + pN_SDE2
  pT_SDE_tt = pT_SDE1 + pT_SDE2

Else
  pN_SDE1 = 0: pN_SDE2 = 0: pN_SDE_tt = 0: pN_SDE_simp1 = 0
  pT_SDE1 = 0: pT_SDE2 = 0: pT_SDE_tt = 0: pT_SDE_simp1 = 0
End If

'VISCO (rappel on calcule les tension avec u négative)
If u < -u0 Then
     If Abs(u) < u2 Then
         pT_vi = pente1 * (Abs(u) - u0)
     Else
         pT_vi = pente2 * (Abs(u) - u2) + pente1 * (u2 - u0)
     End If
Else
  pT_vi = 0
End If


End Sub
```

#### Sub AA_FVtouch_stocke_fm(iX As Integer)

```
zfm = iX
Select Case zfm
  Case Is <= 3 'Edaman 76
   FVto.Seek "=", zfm, -1: If FVto.NoMatch Then Stop
   N_hs = FVto("n_hs")
   FVto.Seek "=", zfm, 1: If FVto.NoMatch Then Stop
   Do While FVto("fm") = zfm
      FVto.Edit
        FVto("u") = FVto("V") * 1000 / N_hs
        FVto.Update
      FVto.MoveNext
  Loop

 Case 4 'Edaman 77
   FVto.Seek "=", zfm, -1: If FVto.NoMatch Then Stop
   N_hs = FVto("n_hs")
   L0 = FVto("L0_fm")
   echY = FVto("Yvrai") / FVto("Ymm")
   FVto.Seek "=", zfm, 1: If FVto.NoMatch Then Stop
   Do While FVto("fm") = zfm
      FVto.Edit
      ' "V": ymi = -u * N_hs / 1000: Stop
    ' "V_L_s": ymi = -u * N_hs / (L0 * 1000)
         FVto("V_L_s") = FVto("Ymm") * echY
        FVto("V") = FVto("V_L_s") * L0
        FVto("u") = FVto("V_L_s") * L0 * 1000 / N_hs
        FVto.Update
      FVto.MoveNext
  Loop

  Case Is <= 13  'Edman 88
   FVto.Seek "=", zfm, -1: If FVto.NoMatch Then Stop
    Select Case zfm
    Case 5, 6: L0 = 6.3 'mmm
    Case 7: L0 = 0.6 'mm
    Case 8, 9, 10, 11: L0 = 8.35 'mm
    Case 12, 13: L0 = 5.5 'mm
    Case Else: Stop
    End Select
   N_hs = L0 * 1000 / 1.05 'on calcule par rapport à la longuer de ref L0__s=2.1 µm
   FVto.Edit
      FVto("L0_fm") = L0
      FVto("N_hs") = Int(N_hs)
   FVto.Update
    ' "V": ymi = -u * N_hs / 1000: Stop
    ' "V_L_s": ymi = -u * N_hs / (L0 * 1000)

   FVto.Seek "=", zfm, 1: If FVto.NoMatch Then Stop
   Do While FVto("fm") = zfm
      FVto.Edit
        FVto("V") = FVto("V_L_s") * L0
        FVto("u") = FVto("V_L_s") * L0 * 1000 / N_hs
        FVto.Update
      FVto.MoveNext
  Loop
```

```
Case 14, 15, 16, 17 'Piazzesi 2002 JPL
    FVto.Seek "=", zfm, -1: If FVto.NoMatch Then Stop
    L0 = IIf(zfm = 14, 5, 6) 'mm
    N_hs = L0 * 1000 / 1.05 'on calcule par rapport à la longuer de ref L0__s=2.1 µm
    xM = IIf(zfm = 14, 6.1, 7.6)
    FVto.Edit
      FVto("L0_fm") = L0
      FVto("S_fm") = xM
      FVto("N_hs") = Int(N_hs)
   FVto.Update

    'verif
   FVto.Seek "=", zfm, -1: If FVto.NoMatch Then Stop
   echX = FVto("Xvrai") / FVto("Xmm")
   echY = FVto("Yvrai") / FVto("Ymm")
   T0m1 = FVto("T_kPa")
   N_hs = FVto("N_hs")
   L0 = FVto("L0_fm")
   xM = FVto("S_fm") 'surface
   FVto.Seek "=", zfm, 1: If FVto.NoMatch Then Stop
   Do While FVto("fm") = zfm
      FVto.Edit
         FVto("T_kPa") = FVto("Xmm") * echX
         FVto("T_Nmm2") = FVto("Xmm") * echX / 1000
         FVto("T_mN") = FVto("Xmm") * xM * echX / 1000
         FVto("pT") = FVto("Xmm") * echX / T0m1
         FVto("u") = FVto("ymm") * echY
         FVto("V") = FVto("ymm") * echY * N_hs / 1000
         FVto("V_L_s") = FVto("ymm") * L0 * echY * N_hs / 1000
      FVto.Update
   FVto.MoveNext: If FVto.EOF Then Exit Do
  Loop

Case 18 'Ford 1973 oct nov dec
    FVto.Seek "=", zfm, -1: If FVto.NoMatch Then Stop
    L0 = 6
    N_hs = L0 * 1000 / 1.1 'on calcule par rapport à la longuer de ref L0__s=2.1 µm
    T0m1 = 285
    FVto.Edit
      FVto("L0_fm") = L0
      FVto("T_kPa") = T0m1
      FVto("N_hs") = Int(N_hs)
   FVto.Update

   'verif
   FVto.Seek "=", zfm, -1: If FVto.NoMatch Then Stop
   echX = FVto("Xvrai") / FVto("Xmm")
   echY = FVto("Yvrai") / FVto("Ymm")
   T0m1 = FVto("T_kPa")
   N_hs = FVto("N_hs")
   L0 = FVto("L0_fm")
   'xM = FVto("S_fm")
   FVto.Seek "=", zfm, 1: If FVto.NoMatch Then Stop
   Do While FVto("fm") = zfm
      FVto.Edit
         FVto("T_kPa") = FVto("Xmm") * echX * T0m1
         FVto("T_Nmm2") = FVto("Xmm") * echX * T0m1 / 1000
         'FVto("T_mN") = FVto("Xmm") * xM * echX / 1000
```

```
         FVto("pT") = FVto("Xmm") * echX
         FVto("u") = FVto("ymm") * echY
         FVto("V") = FVto("ymm") * echY * N_hs / 1000
         FVto("V_L_s") = FVto("ymm") * L0 * echY * N_hs / 1000
      FVto.Update
   FVto.MoveNext: If FVto.EOF Then Exit Do
  Loop

' ****************** ICI ************************

Case 19, 20, 21, 22, 23, 24 ', 17 'Piazzesi 2002 JPL
     FVto.Seek "=", zfm, -1: If FVto.NoMatch Then Stop
     L0 = IIf(zfm <= 21, 15.6, 17.6) 'mm
     T0m1 = Choose(zfm - 18, 165, 198, 209, 158, 188, 198)
     N_hs = L0 * 1000 / 1.25 'on calcule par rapport à la longuer de ref L0__s=2.5 µm
     FVto.Edit
       FVto("L0_fm") = L0
       FVto("T_kPa") = T0m1
       FVto("N_hs") = Int(N_hs)
    FVto.Update

    'verif
   FVto.Seek "=", zfm, -1: If FVto.NoMatch Then Stop
   echX = FVto("Xvrai") / FVto("Xmm")
   echY = FVto("Yvrai") / FVto("Ymm")
   T0m1 = FVto("T_kPa")
   N_hs = FVto("N_hs")
   L0 = FVto("L0_fm")

   FVto.Seek "=", zfm, 1: If FVto.NoMatch Then Stop
   Do While FVto("fm") = zfm
      FVto.Edit
         FVto("pT") = FVto("Xmm") * echX
         FVto("T_kPa") = T0m1 * FVto("Xmm") * echX
         FVto("T_Nmm2") = T0m1 * FVto("Xmm") * echX / 1000
         'FVto("T_mN") = FVto("Xmm") * xM * echX / 1000

           ' "V": ymi = -u * N_hs / 1000: Stop
           ' "V_L_s": ymi = -u * N_hs / (L0 * 1000)
         FVto("V") = FVto("ymm") * echY
         FVto("V_L_s") = FVto("ymm") * echY / L0
         FVto("u") = FVto("ymm") * echY * 1000 / N_hs

      FVto.Update
   FVto.MoveNext: If FVto.EOF Then Exit Do
  Loop

Case 115 'Edaman 88 6mm revu
   FVto.Seek "=", zfm, -1: If FVto.NoMatch Then Stop
   T0m1 = FVto("T_Nmm2")
   FVto.Seek "=", zfm, 1: If FVto.NoMatch Then Stop
   Do While FVto("fm") = zfm
     FVto.Edit
       FVto("pT") = FVto("T_Nmm2") / T0m1
      FVto.Update
     FVto.MoveNext
  Loop
```

```
  Case 25, 26 'Edman88 R1 R1.44
    FVto.Seek "=", zfm, -1: If FVto.NoMatch Then Stop
    L0 = 6.3 'mm
    N_hs = L0 * 1000 / 1.05 'on calcule par rapport à la longuer de ref L0__s=2.1 µm
    xM = 2.5 'IIf(zfm = 14, 6.1, 7.6)
    FVto.Edit
      FVto("L0_fm") = L0
      FVto("S_fm") = xM
      FVto("N_hs") = Int(N_hs)
   FVto.Update
  FVto.Seek "=", zfm, -1: If FVto.NoMatch Then Stop
  echX = FVto("Xvrai") / FVto("Xmm")
  echY = FVto("Yvrai") / FVto("Ymm")
  T0m1 = FVto("T_Nmm2")
  N_hs = FVto("N_hs")
  L0 = FVto("L0_fm")
  xM = FVto("S_fm") 'surface
  FVto.Seek "=", zfm, 1: If FVto.NoMatch Then Stop
  Do While FVto("fm") = zfm
      FVto.Edit
         FVto("T_Nmm2") = FVto("Xmm") * echX
         FVto("T_kPa") = FVto("Xmm") * echX * 1000
         FVto("T_mN") = FVto("Xmm") * xM * echX / 1000
         FVto("pT") = FVto("Xmm") * echX / T0m1
         FVto("V_L_s") = FVto("ymm") * echY
         FVto("V") = FVto("ymm") * echY * L0
         FVto("u") = FVto("ymm") * echY * L0 * 1000 / N_hs
      FVto.Update
  FVto.MoveNext: If FVto.EOF Then Exit Do
 Loop

 Case 29 'Edaman 88 fig 1C 80% T0
  FVto.Seek "=", zfm, -1: If FVto.NoMatch Then Stop
  N_hs = FVto("n_hs")
  L0 = FVto("L0_fm")
  T0m1 = FVto("T_kPa") '163
 ' xM = FVto("S_fm") 'surface
  echX = FVto("Xvrai") / FVto("Xmm")
  echY = FVto("Yvrai") * 1.932 / FVto("Ymm")
  FVto.Seek "=", zfm, 1: If FVto.NoMatch Then Stop
  Do While FVto("fm") = zfm
     FVto.Edit
       FVto("pT") = 0.8 + FVto("Xmm") * echX
       FVto("T_kPa") = FVto("pT") * T0m1
       FVto("T_Nmm2") = FVto("T_kPa") / 1000 'FVto("T_kPa") = FVto("T_Nmm2") * 1000
       'FVto("T_mN") = FVto("T_kPa") * xM / 1000
       FVto("V_L_s") = FVto("Ymm") * echY
       FVto("V") = FVto("V_L_s") * L0
       FVto("u") = FVto("V_L_s") * L0 * 1000 / N_hs
       FVto.Update
     FVto.MoveNext: If FVto.EOF Then Exit Do
 Loop
 Case Else: Stop 'pas BON N°
End Select
End Sub
```
